# Effects of fasting on heat-stressed broiler chickens: part I- growth performance, meat quality, gut histomorphological and microbial responses

**DOI:** 10.1101/2024.09.08.611910

**Authors:** Tanvir Ahmed, Md. Abul Hashem, Afifa Afrin, Ankon Lahiry, Shahina Rahman, Takashi Bungo, Shubash Chandra Das

## Abstract

The current study aimed to optimize the fasting duration in order to mitigate the detrimental effects of heat stress on broilers raised in hot and humid climatic environments. A total of 500 broiler DOCs were assigned to five distinct treatment groups: Tₒ= Non-fasted controlled temperature (24-26℃) (NF-CT), T_1_= Non-fasted heat stressed (30-38℃) (NF-HS), T_2_= 6 hours fasted heat stressed (6-h FHS), T_3_= 8 hours fasted heat stressed (8-h FHS), and T_4_= 10 hours fasted heat stressed (10-h FHS). Each treatment was replicated five times, with 20 birds in each replicate group. As expected, the birds in NF-CT group showed significantly better performances for all the growth parameters, although birds who fasted for 8-h under heat stress exerted better growth and FCR in comparison to the other HS groups. Fasting of birds under heat stress significantly showed the lowest mortality. Like the NF-CT group, birds in 8-h FHS achieved significantly higher dressing percentage, breast meat, liver yields, and the lowest abdominal fat. Fasting for 8- and 10-h significantly increased breast meat pH and water holding capacity and thus reduced cooking loss. Fasting also improved the breast meat color quality by increasing redness (a*) and reducing the hue angle values comparable with the NF-CT group. A significantly upward trend in villi height (VH), width (VW) and crypt depth (CD) of gut segments was also observed in the birds of the 8-h FHS group. Total bacterial and coliform counts in cecum contents were reduced significantly with the increase in the fasting period. Benefit-cost analysis showed better profitability in the 8-h FHS group than other HS groups. Taken altogether, it can be concluded that broiler chicken exposed to 8-h fasting period is an effective approach to mitigate heat stress under hot and humid climatic conditions.

## Introduction

Heat stress, also known as hyperthermia, arises when the birds and animals are unable to regulate their body temperature and produce or absorb more heat than they can dissipate [1]. A devastating and long-lasting ‘heatwave’ observed in recent years during summer engulfed several regions worldwide, including Bangladesh, where the temperature reached almost 43℃ (severe heatwave) [2] that has had a substantial impact on global animal production, notably the production of chickens by accelerating the intensity of heat stress [3–4]. Consequently, a substantial quantity of broiler chickens is dying daily as a result of heatstroke, especially in areas where the birds are being impacted by an intense heatwave [5–6] and has led to a significant crisis in poultry farming. In order to achieve maximum productivity and genetic potential, broiler chickens should be raised in a thermal environment that maintains a temperature range of 18-24°C. Within this range, the birds are able to maintain their inner body temperature at a normal level of 40-41°C [7–9]. However, when the environmental temperature increases due to global warming or climate change, birds lose the ability to regulate body temperature because of excess metabolic heat production, lack of sweat glands, and highly insulated feather covering [10] This leads to a significant increase in body temperature, which can even be lethal to birds if it rises approximately 4°C beyond their normal body temperature [3,11].

Most of the small and medium-scale farmers raise broilers in open-sided houses, exposing the birds to extremely hot and humid natural weather conditions, which has inimical impacts on production performance and even may cause bird mortality due to extreme heat stress (HS) [12–13]. HS has detrimental consequences on various economically significant traits of broilers, such as body weight, weight gain and FCR. HS also has a negative impact on carcass yields and meat quality, leading to a reduction in the proportion of carcass, specifically breast and thigh meat, meat pH levels, shear force, redness and lightness values of thigh and breast meat [14–15]. In addition, HS leads to an augmentation in abdominal adiposity, which further detrimentally affects the visual presentation of the carcass and consumers’ preference for the product [16–17]. Several studies revealed that HS alters the normal histo-morphometry in the digestive tract of chickens, resulting in a decreased area of epithelial cells where nutrients are absorbed [18–21]. The intestinal microbiota is also highly susceptible to HS, with some specific alterations reporting lower levels of beneficial bacteria *(Lactobacillus*, *Bifidobacterium* etc.) and higher levels of harmful bacteria (*Clostridium*, *Coliforms* etc.) [18,19,22,23,24]. The poultry scientists, government sector, environmental professionals, and small-scale farmers are deeply concerned about the HS issue due to its direct impact on the profitability and sustainability of poultry production.

It highlighted the urgent need for climate-resilient farming practices and renewed emphasis on sustainable agriculture to ensure the stability and continuity of world poultry production in the face of ongoing global warming challenges. Therefore, minimizing the adverse effect of HS on the production performance and meat quality of broiler chickens is now a global issue that requires greater attention. Among all attributable factors of HS, metabolic heat production, which is directly related to feed intake, digestion, and absorption in birds, has a substantial impact on production performance, physiology, and meat quality [25]. Numerous approaches are being proposed to combat the HS challenges, the most prominent of which are feed restriction, feed withdrawal, intermittent feeding, and other nutritional management that have drawn much more attention. Feed restriction in broilers, both quantitative and feeding time restrictions, has been employed as one of the most common and effective management practices to combat heat stress challenges in extremely hot and humid climatic conditions [26]. However, in reality, employing a feeding time restriction is much simpler and less stressful than using a quantitative feed restriction [25,27,28], and also confers a more beneficial impact on physiology and behavior of meat-type birds [29]. Feeding time restriction is a feeding strategy in which birds are not given access to feed for a fixed or alternative duration of time, primarily practiced during the second half of the raising period, from day 21 onward. The benefits of feed withdrawal or fasting approaches include decreased metabolic heat production from food digestion, absorption, and metabolism, as well as reduced body temperature and mortality during the hottest part of the day. However, prolonged fasting may decrease the birds’ weight gain [30]. Fasting can also induce metabolic acidosis to counterbalance the respiratory alkalosis associated with mortality and affect the response of fasted birds exposed to heat [31].

Without comprehending the scientific rationale, broiler farmers in many tropical countries, including Bangladesh, have been practicing fasting regimes during the hottest part of the day for different durations. They are still dubious about the optimal fasting duration that broiler chickens must fast in order to recover from the adverse effects of heat stress. Additionally, it is important to determine which of the fasting durations have a significant impact on the birds’ growth performance, meat quality, gut health, gut microbiota and other relevant metrics. Several studies have investigated the effect of feed withdrawal or fasting strategies on the performance of broiler chickens. However, there is scanty information regarding the optimal duration of fasting that would reduce heat stress in broilers while enhancing growth, meat quality, gut health, and reducing the pathogenic microbial load. Therefore, this study aimed to determine the most effective duration of fasting to mitigate the heat stress challenge on broiler production by focusing on the impact of growth performance, meat quality, gut histomorphology, and microbiological reactions in broiler chickens.

## Materials and methods

### Experimental site and ethical statement

The experiment was conducted during the hot summer at Bangladesh Agricultural University Poultry Farm, Bangladesh. All operational activities related to the bird’s raising, slaughtering, overall management, and sample collection were approved by the Animal Welfare and Experimentation Ethics Committee (AWEEC) of Bangladesh Agricultural University (Approval number: AWEEC/BAU/2024-27).

### Experimental design, chick collection and distribution

A total of 500-day-old commercial broiler chicks were selected for the experimental trial and randomly assigned to five distinct treatment groups with five replications and 20 birds for each replication. From day one, birds were distributed in accordance with the completely randomized design (CRD) to the treatment groups: namely, T_ₒ_= Non-fasted control temperature (24-26°C) (NF-CT), T_1_= Non-fasted heat stressed (30-38°C) (NF-HS), T_2_= 6 hours fasted heat stressed (6-h FHS), T_3_= 8 hours fasted heat stressed (8-h FHS) and T_4_= 10 hours fasted heat stressed (10-h FHS). The broiler chicks were collected from a reputed hatchery of AG Agro Industries Limited, Gazipur, Bangladesh. Immediately upon arrival, the DOCs were weighed and then randomly distributed to floor pens as per treatments and replications. For the first three weeks, the birds in all treatment groups were provided feed and water *ad-libitum* and kept under identical care and management; after that, different feeding management approaches were employed.

### House preparation, brooding and other management practices

The experimental birds were reared in two separate houses: first, in a gable-type open-sided poultry house where the birds of four treatment groups (NF-HS, 6-h FHS, 8-h FHS and 10-h FHS) were exposed to high ambient temperatures ranging from 30-38°C; and second, in an environmentally controlled chamber (NF-CT) with temperatures ranging from 24-26°C. Each pen was 22 square feet, predicting about 18 kg live weight per square meter (around 1.1 square feet per bird), where the birds were reared for five weeks. The experimental house was well-cleaned, scrubbed, and disinfected and finally left to rest for seven days before the arrival of the DOCs. Before the DOCs being placed in each pen, all necessary equipment was carefully cleaned, sanitized, and disinfected. Fresh and dry rice husk was strewn over the floor to a depth of 3 cm as bedding materials. Just before the arrival of DOCs, the newspaper was laid over the litter materials as chick paper, and the chicks were brooded in respective pens for seven days using two 100-watt electric bulbs. The birds were exposed to 23 hours of lighting for the first three days of the brooding period, and then, they maintained a standard lighting schedule for the remaining periods of rearing, following management guidelines by adjusting both natural and artificial lighting. Some management measures were performed to maintain the recommended temperature and humidity, such as turning off one or both lights, elevating the bulbs, withdrawing curtains, and so on. A total of six automatic thermo-hygrometers (HTC-2, Velveeta, Makkar Trading Company, India) were set at six different points in the house to measure the temperature and humidity of the experimental house. After 14 days of rearing, the newspaper was removed from the litter. Litter was stirred at least twice a day throughout the experimental period.

### Feed and water management

The birds were fed broiler starter crumble feed from day 1 to day 21 and broiler grower pellet feed for the rest of the experimental period. Feed for experimental birds was purchased from a reputed feed mill named Ag Agro Industries Limited, Mauna, Gazipur, Dhaka, was used for the feeding of experimental birds (Table 1). According to the research design, all of the experimental birds were not subjected to treatment-wise fasting approaches for the first three weeks, and therefore, an *ad libitum* feed was provided. Afterward, the feed was delivered at least twice daily in such a way that the birds in the non-fasting groups had access to feed at all times, whereas the feeders in the fasted groups were kept empty for 6, 8, and 10 hours, respectively. The birds in NF-CT and NF-HS groups have access to feed all day long (*ad-libitum*), and for 6-h, 8-h and 10-h FHS groups feeders were kept empty for 6 hours (11 am to 5 pm), 8 hours (10 am to 6 pm) and 10 hours (8 am to 6 pm) respectively.

**Table 1.** Nutrient composition of broiler diets.

| Name of the Nutrients | Amount |  |
| --- | --- | --- |
|  | Starter diet (1-21 days) | Grower diet (22-35 days) |
| ME (kcal/kg) | 3050 | 3150 |
| CP (%) | 22.00 | 21.00 |
| Moisture (%) | 12.00 | 12.00 |
| Crude fat (%) | 6.00 | 7.00 |
| Crude fiber (%) | 4.00 | 6.00 |
| Ash (%) | 8.00 | 8.00 |
| Calcium (%) | 0.98 | 0.98 |
| Av. Phosphorus (%) | 0.75 | 0.75 |

Where, ME= Metabolizable energy; kcal= Kilo calorie; CP=Crude protein; Av.=Available

### Immunization of the birds, biosecurity and sanitation measures

Experimental birds were administered with an IB+ND vaccine (RaniVexTM Plus, which combines the LaSota strain of ND Virus and the Massachusetts B48 strain of IB Virus in live, freeze-dried form) at the age of 4 days, subsequently receiving a booster dose on day 20. Furthermore, on day 10, the IBD vaccine (GumboMedTM Vet, including live attenuated IBD or Gumboro virus intermediate strain) was administered, followed by a booster dose on day 17. All of the birds were inoculated with eye drops. A stringent biosecurity protocol was implemented throughout the study area, and outsider access to the experimental shed was strictly prohibited. Both hands and feet were thoroughly cleansed and sprayed with the Virkon S disinfectant (DuPontTM Virkon® S, Antec International, USA) to avoid contamination. Separate footwear and aprons were also utilized before entering the experimental unit as part of hygiene and sanitation, and a foot wash containing potassium permanganate was installed at the entrance to the experimental shed. To ensure strict biosecurity controls, the feeders and drinkers were cleaned and sanitized daily, and the dead birds, if any, were appropriately disposed of.

### Carcass yield and meat quality determination

Following the completion of the experimental trial, two birds of average weight were randomly chosen from each treatment group to assess carcass yield. Feed withdrawal was performed around 12 hours before the birds were slaughtered to ensure that their crops were almost empty. However, water was consistently provided to ensure normal bleeding. Subsequently, the birds were weighed and anesthetized with ethanol before slaughtering ethically according to the Animal Welfare Act and processed by following the standard protocol. To determine the dressing percentage of experimental birds, the carcass weight was first calculated by subtracting the weight of feathers, skin, visceral organs, and other inedible components from the live weight of the birds and then expressed as a percentage of the live weight of the birds. The weight of various parts of the experimental birds, including the thigh, drumstick, breast meat, wing, head, liver, gizzard, heart, neck, shank, and abdominal fat, were determined using a digital balance. The organ weights were determined by calculating the percentage of each organ’s weight in relation to the total live weight of the birds. Following the processing, the breast meat samples were collected and held at a temperature of 4℃ to measure meat quality parameters. The pH of the breast meat was determined 24 hours after slaughtering by inserting the probe of a digital HI 99163 pH meter (manufactured by HANNA Instruments. Inc. in Highland Industrial Park, USA) into the breast meat sample. The color of breast meat samples was assessed on the surface region using a Konica Choma Metres CR-410 (Konica Minolta Inc., Tokyo, Japan) and quantified using the CIE LAB system, which measures the characteristics of lightness (L*), redness (a*), and yellowness (b*). The hue angle and saturation index (SI) were determined using the formulas: “Hue angle = tan^-1^ (b*/a*)” and “Saturation index (SI) = (a*^2^+b*^2^)^1/2^” [32–33]. The water holding capacity (WHC) was determined using a centrifugation assay. About 1 g breast meat sample from each replication was cut into cubes and centrifuged for 10 minutes at 10,000 rpm. The WHC was determined by quantifying the water exudation from the meat sample through centrifugation, and the results were expressed as a percentage. The breast meat sample weighing approximately 15 g was first weighed and then placed in a double-layer polythene bag with an identifying mark. Then, it was submerged in a water bath set at 80°C for 30 minutes to measure cooking loss. Following the cooking process, the samples were subjected to a 10-minute air drying, after which their weight was measured again in order to determine the percentage of cooking loss. For the determination of drip loss, a consistent amount of approximately 25 g (wet weight) of meat with a regular shape was cut from the breast at the same position for each sample and weighed. The sample was thereafter enclosed in a hermetically sealed container by suspending it from a string and stored in a refrigerator maintained at 4°C. After 24 hours, samples were extracted from the freezer and subsequently weighed to determine the percentage of drip loss.

### Gut sample collection and measurement of histomorphometrics

After the dissection of sampling broilers, sections of the duodenum, jejunum, and ileum measuring 1.0-1.5 cm in length were obtained. These sections were then rinsed with phosphate-buffered saline (PBS) to remove the intestinal content. For histological analysis, the duodenum segment was taken from the midpoint of the duodenal loop, the jejunum segment from the location midway between the end of the duodenal loop and Meckel’s diverticulum, the ileum segment was taken from merely five inches behind the ileocecal junction. Collected segments were then fixed in a neutral-buffered formalin (10%) solution. The tissue samples were subsequently dehydrated using increasing concentrations of ethanol (Merk, Damstadt, Germany), cleared using xylene (Merck, Damstadt, Germany), and finally embedded in paraffin. Tissue sections measuring 5 µm in thickness were obtained using a rotary microtome (model RM2016, LEIC, Germany), which were then placed on glass slides, with each slide numbered to maintain the order of the sections. The sections were subsequently stained with hematoxylin and eosin to facilitate histomorphometric analysis. Finally, the villus height, width and crypt depth were determined using a light microscope with an image analyzer (Image-ProPlus 6.0) and ImageJ software (Version 1.8.0). Assessment of villus height (VH) and width (VW) were obtained from at least five full villi, as well as the depth of crypt (CD) was measured from five randomly chosen sites of the crypt from each intestinal sample and then the average value was taken in order to improve the accuracy of evaluating the gut health of birds.

### Cecal bacterial count

At the end of the experiment, three birds from each replicate group were chosen randomly and euthanized humanely. Their intestines were then dissected to determine the bacterial count. The cecum was extracted from each dissected bird using a sterile pair of scissors and compressed with sterile forceps to obtain approximately 1 g of fresh cecal contents, which were then collected in a sterile falcon tube. A vortex was used to thoroughly mix about 1 g of fresh cecal materials and 10 ml of PBS. Subsequently, a tenfold serial dilution was performed inside the Eppendorf tube, resulting in a total of 10^6^ dilutions. Subsequently, 50 μl aliquots from every dilution were placed in petri dishes containing distinct media. Then, the diluted samples were evenly distributed onto bacteriological media, including Nutrient agar and MacConkey agar for *E. coli* count and Blood agar for counting total (anaerobic) bacteria. The Nutrient agar and MacConkey agar plates were inoculated and then placed in an incubator at 37°C overnight, whereas Blood agar plates were incubated anaerobically at 42°C. Then, the bacterial colony was enumerated and subsequently subjected to morphological analysis using Gram’s staining. The manually measured bacterial colonies were multiplied by the dilution factor and volume factor to determine the CFU (Colony Forming Unit) of anaerobic bacteria and *E. coli*. The results were expressed as Log_10_ CFU/g of cecal content.

### Data collection and record keeping

The temperature and humidity of the study area were recorded six times daily (at 6:00 am, 10:00 am, 12:00 pm, 2:00 pm, 5:00 pm, and 10:00 pm). The temperature humidity index (THI) was calculated according to the formula of Marai et al. [34] and Hahn et al. [35]. The initial body weight and feed supply of the birds were recorded on arrival and on a weekly basis during the experimental period. Every week, the birds’ final body weight was subtracted from their initial body weight to determine their body weight gain. Residual feed was recorded weekly to determine the weekly feed intake per bird. Subsequently, by dividing the recorded weekly feed intake by the weight gain of experimental birds, the feed conversion ratio (FCR) was computed. The mortality was recorded daily to determine their survivability. Data on carcass yield, meat quality, gut health and cecal bacterial load was recorded and measured at the end of the experiment.

### Calculation of production cost and profit

The cost-benefit analysis was conducted by considering the expenditures associated with DOCs, feed, feeder, drinker, litter, medicines, and other related expenses. The total cost (TC) was subtracted from the total revenue (TR) to determine profit.

### Statistical analysis

The recorded and calculated data obtained during the experiment were analyzed using the One-Way Analysis of Variance (ANOVA) technique. This analysis was performed using the SPSS, Statistical Computer Package Program 20.00 [36], and followed the principles of Completely Randomized Design (CRD). The Standard Error (SE) was computed to assess significant variations among treatments in cases where ANOVA indicated significant differences. Duncan’s Multiple Range Test (DMRT) was employed to compare the significance among various treatments.

## Results

### Temperature humidity index (THI)

The temperature humidity index (THI) observed during the last two weeks of the experiment is illustrated in Fig 1. The results indicated that birds raised in hot and humid environmental conditions (NF-CT, 6-h FHS, 8-h FHS and 10-h FHS) experienced a THI ranging from 84 to 89. In contrast, birds raised in a controlled environment faced a THI ranging from 73 to 74 during the 3^rd^ to 5^th^ weeks of age.

**Fig 1.**
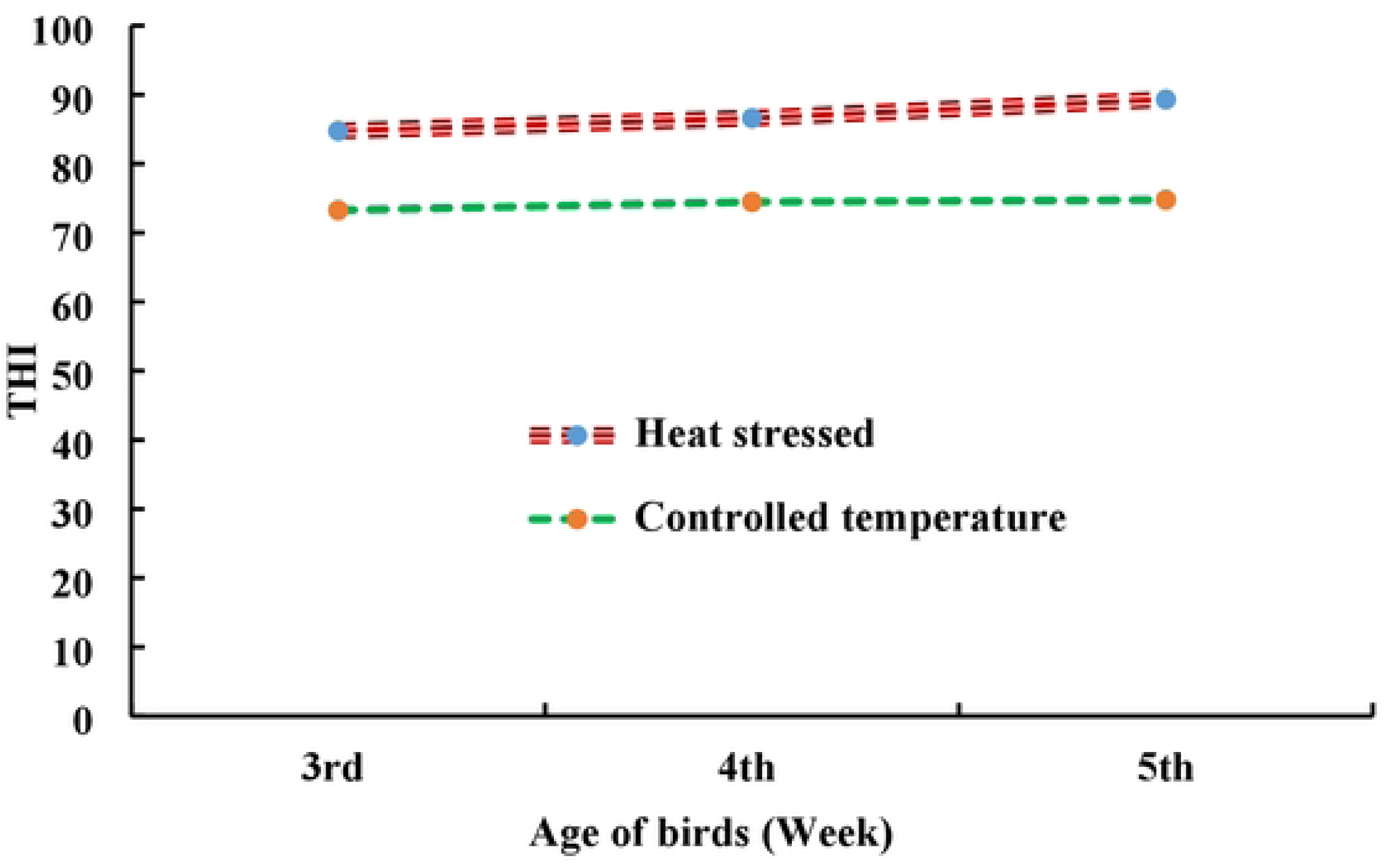
THI during the experimental period for both controlled temperature and heat­stressed groups. Where, THI= Temperature humidity index.

### Growth performance and mortality

Table 2 illustrates the impact of fasting on the growth performance of broilers raised in hot and humid climates. The results indicated that there were no significant differences in cumulative growth performance among the treatment groups until 3^rd^ week when the birds had not yet been subjected to fasting. However, at the end of the experimental trial (after 5 weeks), the NF-CT group demonstrated a statistically significant increase in body weight (BW) and body weight gain (BWG) compared to the NF-HS and other FHS groups (p=0.00). Within the heat-stressed groups, no statistical differences were observed in BW and BWG, although the birds that fasted for 6-h and 8-h under heat stress showed numerically better BW and BWG at 5^th^ weeks of age. As anticipated, the feed intake (FI) was considerably greater (p=0.03) in the NF-CT group (3186.79 g/b) compared to the birds in the non-fasting (2974.68 g/b) or 8-h fasting (3001.00 g/b) groups under heat stress conditions. The remaining two treatment groups, subjected to heat stress and fasting for 6-h and 10-h, consumed a moderate quantity of feed. In addition to growth performance, the birds in the NF-CT group exhibited a significantly improved feed conversion ratio (FCR=1.49) compared to the other four treatment groups. Among the heat-stressed groups, the birds that were subjected to an 8-h fasting showed considerably improved feed conversion ratio (FCR=1.60); however, relatively greater FCR values were seen in the non-fasting heat-stressed group (NF-HS=1.63), as well as in the birds with 6-h (1.63) and 10-h (1.68) fasting periods. The birds reared in non-fasting under heat stress had a significantly higher mortality rate (6%). However, no birds that underwent 8-h and 10-h fasting or were kept in a controlled environment (NF-CT group) died throughout the experimental period.

**Table 2.** Growth performance and mortality of broilers subjected to different periods of fasting under controlled and high ambient temperature.

| Parameters | Age (Day) | NF-CT | NF-HS | 6-h FHS | 8-h FHS | 10-h FHS | p-value | LS |
| --- | --- | --- | --- | --- | --- | --- | --- | --- |
| Initial wt. (g/b) | D: 0 | 43.86±0.47 | 43.10±0.20 | 43.42±0.30 | 43.34±0.29 | 43.28±0.29 | 0.76 | NS |
| FBW (g/b) | D: 1-21 | 1028.00±18.18 | 960.16±13.53 | 957.55±19.02 | 970.56±22.74 | 954.81±20.27 | 0.06 | NS |
|  | D: 1-35 | 2168.25<sup>a</sup>±30.39 | 1873.96<sup>b</sup>±36.99 | 1919.87<sup>b</sup>±33.06 | 1921.14<sup>b</sup>±45.05 | 1861.14<sup>b</sup>±6.14 | 0.00 | ** |
| BWG (g/b) | D: 1-21 | 984.00±18.18 | 917.06±13.52 | 914.12±18.92 | 927.22±22.88 | 911.53±20.28 | 0.07 | NS |
|  | D: 1-35 | 2126.10<sup>a</sup>±30.43 | 1830.86<sup>b</sup>±37.07 | 1876.45<sup>b</sup>±33.15 | 1877.80<sup>b</sup>±45.19 | 1817.86<sup>b</sup>±6.02 | 0.00 | ** |
| FI (g/b) | D: 1-21 | 1184.08±12.46 | 1142.52±24.11 | 1138.04±19.75 | 1178.10±17.39 | 1139.39±20.76 | 0.29 | NS |
|  | D: 1-35 | 3186.79<sup>a</sup>±40.97 | 2974.68<sup>b</sup>±38.77 | 3064.18<sup>ab</sup>±64.59 | 3001.00<sup>b</sup>±38.02 | 3052.04<sup>ab</sup>±34.96 | 0.03 | * |
| FCR | D: 1-21 | 1.20±0.01 | 1.24±0.01 | 1.24±0.00 | 1.27±0.06 | 1.25±0.08 | 0.10 | NS |
|  | D: 1-35 | 1.49<sup>c</sup>±0.02 | 1.63<sup>ab</sup>±0.02 | 1.63<sup>ab</sup>±0.01 | 1.60<sup>b</sup>±0.02 | 1.68<sup>a</sup>±0.18 | 0.00 | ** |
| Mortality (%) | D: 1-21 | 0.00 | 0.00 | 0.00 | 0.00 | 0.00 | - | - |
|  | D: 1-35 | 0.00<sup>b</sup>±0.00 | 6.00<sup>a</sup>±2.45 | 2.00<sup>ab</sup>±2.00 | 0.00<sup>b</sup>±0.00 | 0.00<sup>b</sup>±0.00 | 0.04 | * |

<sup>a,b,c</sup> Means bearing uncommon superscripts in a row differ significantly ( $P \leq 0.05$ ); NS= Non-significant ( $P > 0.05$ ); \*\* = ( $P < 0.01$ ); \* = ( $P < 0.05$ ); Value indicates – Mean ± Standard Error (SE); LS = Level of significance; NF-CT= Non-fasted controlled temperature; NF- HS= Non-fasted heat stressed; 6-h FHS= 6-hour fasted heat stressed; 8-h FHS= 8-hour fasted heat stressed; 10-h FHS= 10-hour fasted heat stressed; FBW= Final body weight; BWG= Body weight gain; FI= Feed intake; FCR= Feed conversion ratio.

### Impact of fasting on carcass yield

The carcass yields and proportions of various organs in broilers, relative to body weight, are displayed in Table 3. The research findings unequivocally showed that the carcass weight, dressing percentage (DP%), breast meat, liver weight, and abdominal fat percentage were significantly influenced by the various treatment groups. The carcass weight was notably greater in the NF-CT and 8-h FHS groups, followed by the birds subjected to 6-h fasting. The birds with NF-HS or those subjected to 10-h fasting, however, exhibited the lowest carcass weights. The dressing output, beast meat, and liver weight were significantly higher in broilers raised in NF-CT or 6-h FHS compared to all other treatment groups. The percentage of abdominal fat was significantly highest in the NF-CT (1.69 %) and NF-HS (1.62 %) groups but predictably lower in the birds subjected to fasting, with a noticeable trend of gradual decrease as the fasting period increased (1.22 %, 1.04 %, and 0.86 % for 6-h, 8-h, and 10-h FHS groups, respectively).

**Table 3.**
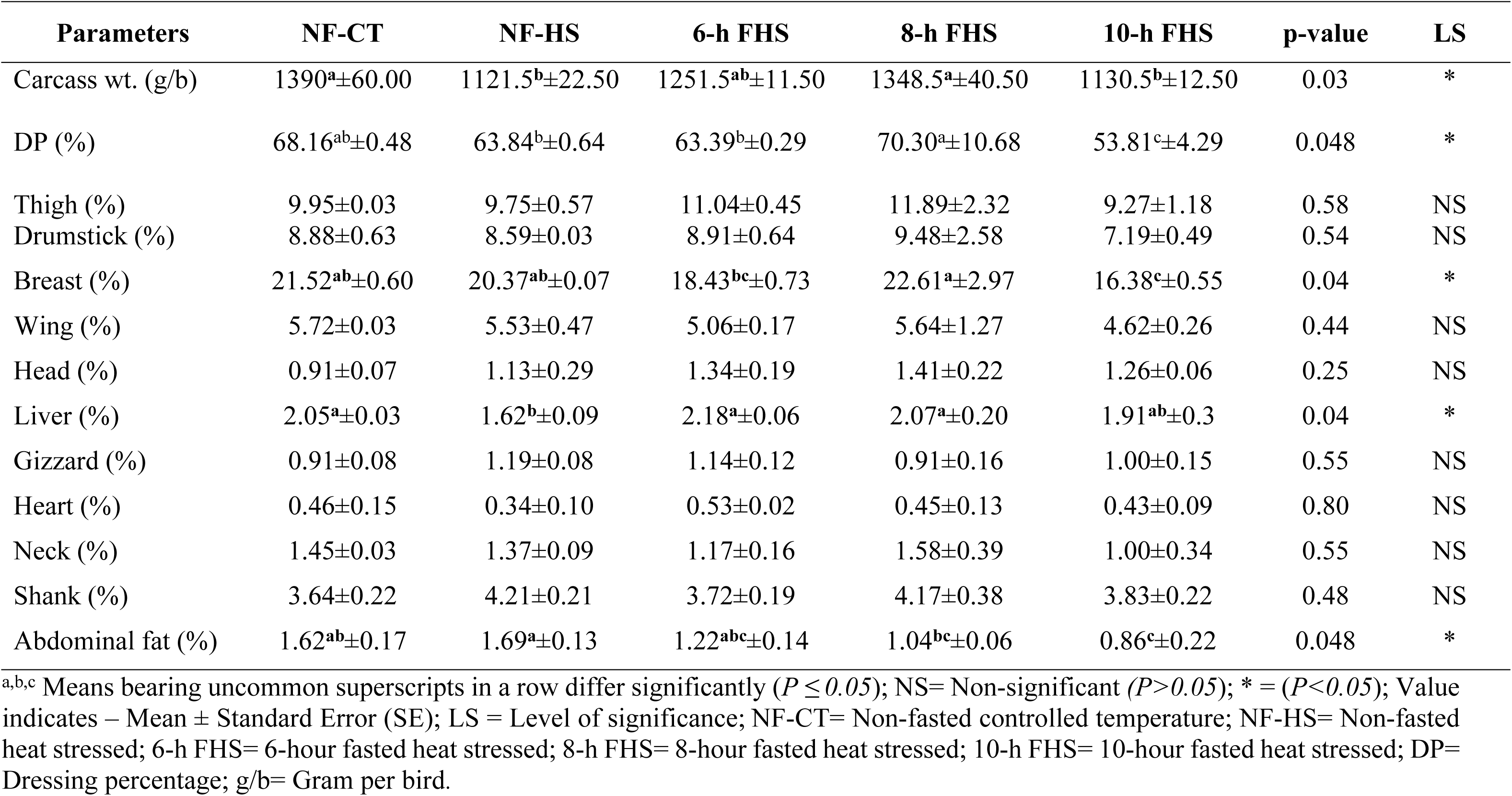
Carcass yields and proportion of other organs of broilers exposed to different fasting under controlled and high ambient temperature.

| Parameters | NF-CT | NF-HS | 6-h FHS | 8-h FHS | 10-h FHS | p-value | LS |
| --- | --- | --- | --- | --- | --- | --- | --- |
| Carcass wt. (g/b) | 1390<sup>a</sup>±60.00 | 1121.5<sup>b</sup>±22.50 | 1251.5<sup>ab</sup>±11.50 | 1348.5<sup>a</sup>±40.50 | 1130.5<sup>b</sup>±12.50 | 0.03 | * |
| DP (%) | 68.16<sup>ab</sup>±0.48 | 63.84<sup>b</sup>±0.64 | 63.39<sup>b</sup>±0.29 | 70.30<sup>a</sup>±10.68 | 53.81<sup>c</sup>±4.29 | 0.048 | * |
| Thigh (%) | 9.95±0.03 | 9.75±0.57 | 11.04±0.45 | 11.89±2.32 | 9.27±1.18 | 0.58 | NS |
| Drumstick (%) | 8.88±0.63 | 8.59±0.03 | 8.91±0.64 | 9.48±2.58 | 7.19±0.49 | 0.54 | NS |
| Breast (%) | 21.52<sup>ab</sup>±0.60 | 20.37<sup>ab</sup>±0.07 | 18.43<sup>bc</sup>±0.73 | 22.61<sup>a</sup>±2.97 | 16.38<sup>c</sup>±0.55 | 0.04 | * |
| Wing (%) | 5.72±0.03 | 5.53±0.47 | 5.06±0.17 | 5.64±1.27 | 4.62±0.26 | 0.44 | NS |
| Head (%) | 0.91±0.07 | 1.13±0.29 | 1.34±0.19 | 1.41±0.22 | 1.26±0.06 | 0.25 | NS |
| Liver (%) | 2.05<sup>a</sup>±0.03 | 1.62<sup>b</sup>±0.09 | 2.18<sup>a</sup>±0.06 | 2.07<sup>a</sup>±0.20 | 1.91<sup>ab</sup>±0.3 | 0.04 | * |
| Gizzard (%) | 0.91±0.08 | 1.19±0.08 | 1.14±0.12 | 0.91±0.16 | 1.00±0.15 | 0.55 | NS |
| Heart (%) | 0.46±0.15 | 0.34±0.10 | 0.53±0.02 | 0.45±0.13 | 0.43±0.09 | 0.80 | NS |
| Neck (%) | 1.45±0.03 | 1.37±0.09 | 1.17±0.16 | 1.58±0.39 | 1.00±0.34 | 0.55 | NS |
| Shank (%) | 3.64±0.22 | 4.21±0.21 | 3.72±0.19 | 4.17±0.38 | 3.83±0.22 | 0.48 | NS |
| Abdominal fat (%) | 1.62<sup>ab</sup>±0.17 | 1.69<sup>a</sup>±0.13 | 1.22<sup>abc</sup>±0.14 | 1.04<sup>bc</sup>±0.06 | 0.86<sup>c</sup>±0.22 | 0.048 | * |

<sup>a,b,c</sup> Means bearing uncommon superscripts in a row differ significantly ( $P \leq 0.05$ ); NS= Non-significant ( $P > 0.05$ ); \* = ( $P < 0.05$ ); Value indicates – Mean ± Standard Error (SE); LS = Level of significance; NF-CT= Non-fasted controlled temperature; NF-HS= Non-fasted heat stressed; 6-h FHS= 6-hour fasted heat stressed; 8-h FHS= 8-hour fasted heat stressed; 10-h FHS= 10-hour fasted heat stressed; DP= Dressing percentage; g/b= Gram per bird.

### Impact of fasting on meat quality

Table 4 illustrates the impact of fasting on the meat quality characteristics of broilers subjected to controlled temperature and hot-humid climatic conditions. The breast meat pH was significantly enhanced in the NF-CT (6.46) and 8-h FHS groups (6.44), whereas the birds fed *ad-libitum* (NF-HS) exhibited the lowest breast meat pH (6.06) in hot and humid weather conditions. The water holding capacity (WHC) was found to be the lowest (90.61%) in the NF-HS group, whereas the 8-h and 10-h FHS groups exhibited the highest WHC values (93.46% and 96.56%, respectively). However, there was a considerably greater cooking loss (p<5.00) seen in the birds of the NF-HS group (33.89%) compared to the others. Conversely, the 8-h FHS group had the lowest cooking loss (21.74%) among all the groups. The drip loss remained unaffected by the different treatments.

**Table 4.** Meat quality of broilers exposed to different periods of fasting under controlled temperature and heat stressed condition.

| Parameters | NF-CT | NF-HS | 6-h FHS | 8-h FHS | 10-h FHS | P-value | LS |
| --- | --- | --- | --- | --- | --- | --- | --- |
| pH | 6.46<sup>a</sup>±0.01 | 6.06<sup>c</sup>±0.03 | 6.30<sup>b</sup>±0.02 | 6.44<sup>a</sup>±0.03 | 6.37<sup>ab</sup>±0.03 | 0.001 | ** |
| WHC | 94.91<sup>ab</sup>±1.70 | 90.61<sup>b</sup>±1.38 | 92.39<sup>ab</sup>±0.90 | 96.56<sup>a</sup>±0.84 | 93.46<sup>a</sup>±0.79 | 0.045 | * |
| CL (%) | 24.97<sup>bc</sup>±1.93 | 33.89<sup>a</sup>±1.43 | 32.11<sup>ab</sup>±0.50 | 21.74<sup>c</sup>±2.51 | 26.19<sup>bc</sup>±2.44 | 0.028 | * |
| DL (%) | 2.29±0.34 | 2.71±0.71 | 2.55±0.04 | 2.26±0.30 | 2.52±0.39 | 0.899 | NS |

<sup>a,b,c</sup> Means bearing uncommon superscripts in a row differ significantly ( $P \leq 0.05$ ); NS= Non-significant ( $P > 0.05$ ); \*\* = ( $P < 0.01$ ); \* = ( $P < 0.05$ ); Value indicates – Mean ± Standard Error (SE); LS = Level of significance; NF-CT= Non-fasted controlled temperature; NF-HS= Non-fasted heat stressed; 6-h FHS= 6-hour fasted heat stressed; 8-h FHS= 8-hour fasted heat stressed; 10-h FHS= 10-hour fasted heat stressed; WHC= water holding capacity; CL=Cooking loss; DL=Drip loss.

When examining the color of the breast meat (Fig 2), fasting of the birds did not have any influence on the lightness (L*) and yellowness (b*), but it did considerably increase redness (a*) in the birds fasted for 8-h (9.03) and 10-h (8.78) as compared to the other groups. The NF-HS group displayed the lowest recorded value of redness (2.82). While the differences in the lightness (L*) value of broiler breast meat among treatment groups were not statistically significant, the NF-HS group exhibited a numerically greater L* value of 53.99 compared to the other treatment groups. Broilers in the NF-HS group exhibited a significantly higher hue angle of 77.43⁰ in their breast meat, while all other treatments showed lower hue angle values ranging from 52-61⁰. Various treatments did not significantly influence the saturation index (SI) of broiler breast meat. The NF-HS group had numerically lowest SI (9.75), followed by the NF-CT, 6-h FHS, and 10-h FHS groups with values ranging from 13 to 15. The highest SI of 18.34 was found in the 8-h FHS group.

**Fig 2.**
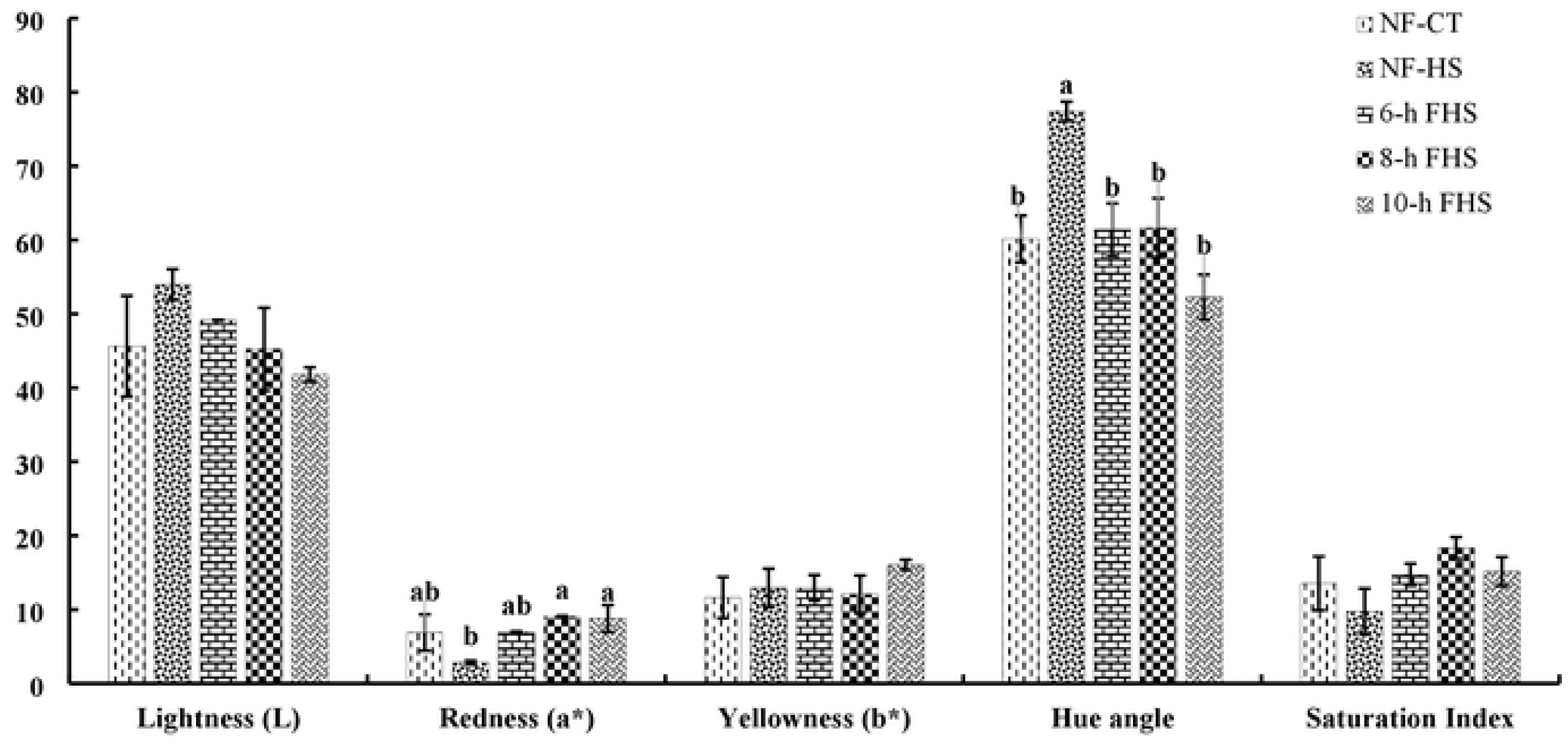
Changes in the meat color of broilers exposed to different period of fasting under controlled and high ambient temperature condition. Where, a,b Means bearing uncommon superscripts in a row differ significantly *(P* :S *0.05);* NS= Non significant *(P>0.05); \*\** = *(P<O.OI); \** = *(P<0.05);* Value indicates - Mean ± Standard Error (SE); LS = Level of significance; NF-CT= Non-fasted controlled temperature; NF-HS= Non-fasted heat stressed; 6-h FHS= 6-hour fasted heat stressed; 8-h FHS= 8-hour fasted heat stressed; 10-h FHS= 10-hour fasted heat stressed.

### Gut histomorphological changes among the experimental birds

Table 5 presents the impact of various fasting approaches on the histomorphological changes in three distinct gut segments (duodenum, jejunum, and ileum) of broiler chickens subjected to controlled temperature and extremely hot-humid weather conditions. The study indicated that the villi height (VH), villi width (VW), and crypt depth (CD) of the duodenum, jejunum, and ileum were considerably higher (p<0.05) in the birds of the NF-CT group compared to the NF-HS group, except for the VW of the ileum, which did not show a significant difference among the treatment groups. Conversely, the ratio of villi height to crypt depth (VH: CD) was significantly greater in the NF-HS group compared to the NF-CT. When examining all groups exposed to heat stress, it is noteworthy that birds subjected to 8-h fasting exhibited significantly higher values for VH, VW, CD, and VH:CD ratio in the duodenum, jejunum, and ileum. These values were statistically similar to, and sometimes even better than, those observed in the NF-CT group. Conversely, the birds in both the NF-HS and 10-h FHS groups had the lowest values for these histological traits. The data clearly indicate that there is a consistent increase in VH, VW, CD, and VH:CD in the gut segments (duodenum, jejunum, and ileum) as the fasting period extends up to 8-h under heat stress. However, it can be noted that fasting for longer than 8-h negatively affects the gut health of commercial broiler chickens.

**Table 5:**
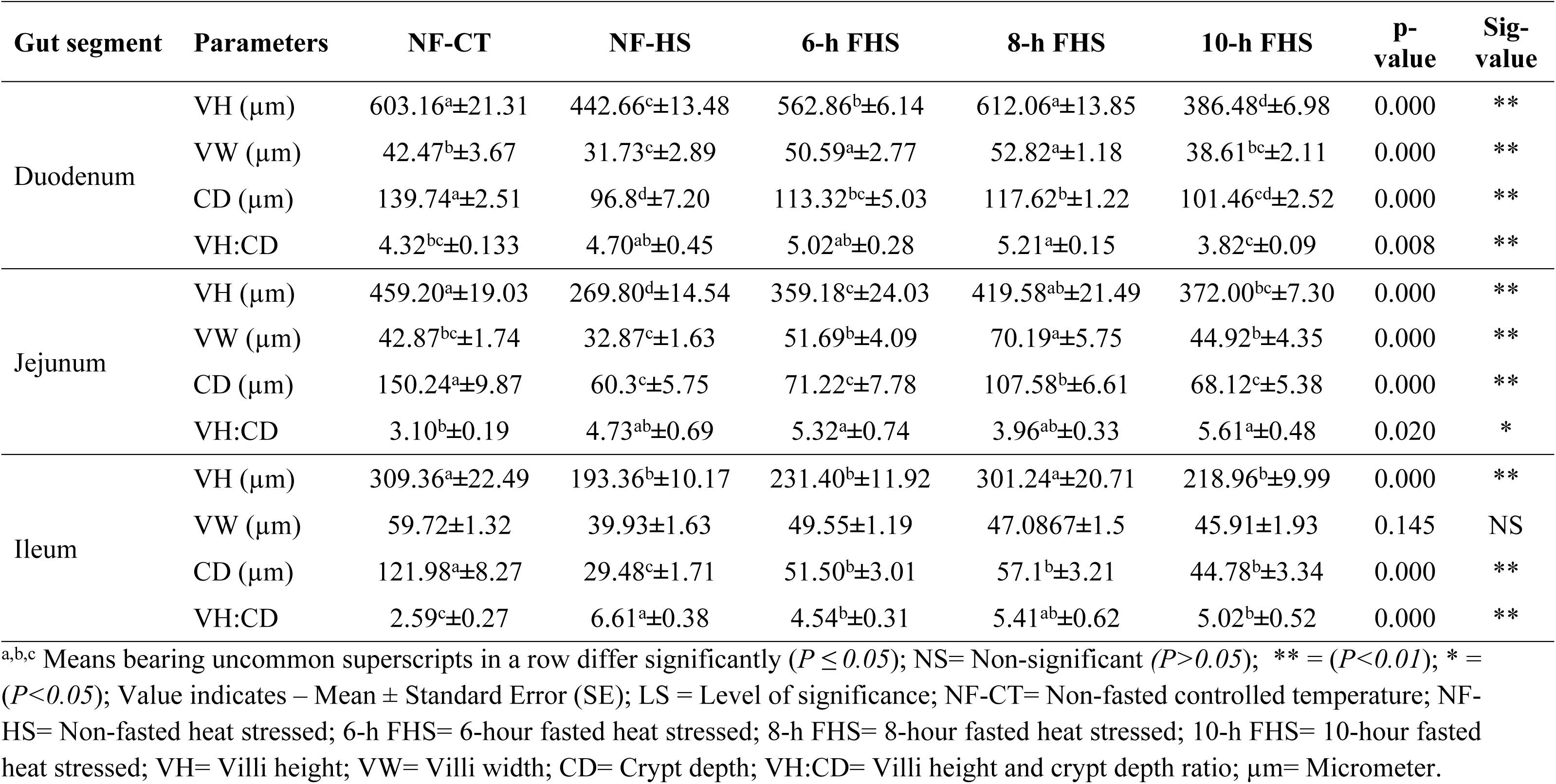
Histomorphological changes of gut segments of broilers affected by fasting approaches under controlled temperature and heat stressed condition.

### Impact of fasting regimes on the bird’s cecal microbial profile

The alterations in the microbial profile of cecum among different treatment groups are displayed in Table 6. The NF-HS group had the highest bacterial count (5.26×10^7^ CFU/g), followed by the 6-h FHS group (3.99×10^7^ CFU/g). The lowest bacterial count, however, was observed in the birds kept either non-fasting in the controlled environment (NF-CT) (2.05×10^7^CFU/g) or fasting for 8-h and 10-h under heat stress (2.27×10^7^ and 1.97×10^7^ CFU/g, respectively). An almost similar pattern was observed with coliform bacteria, with the NF-HS group showing a considerably greater population (2.01×10^7^ CFU/g) and the NF-CT, 8-h, and 10-h FHS groups having the lowest populations (0.91×10^7^, 0.90×10^7^, and 0.85×10^7^ CFU/g, respectively). To summarize, heat stress leads to an increase in both coliforms and total bacterial counts in the ceca of broiler chickens. However, there is a gradual and downward trend of a decrease in the overall bacterial population with an increase in the length of fasting periods.

**Table 6.**
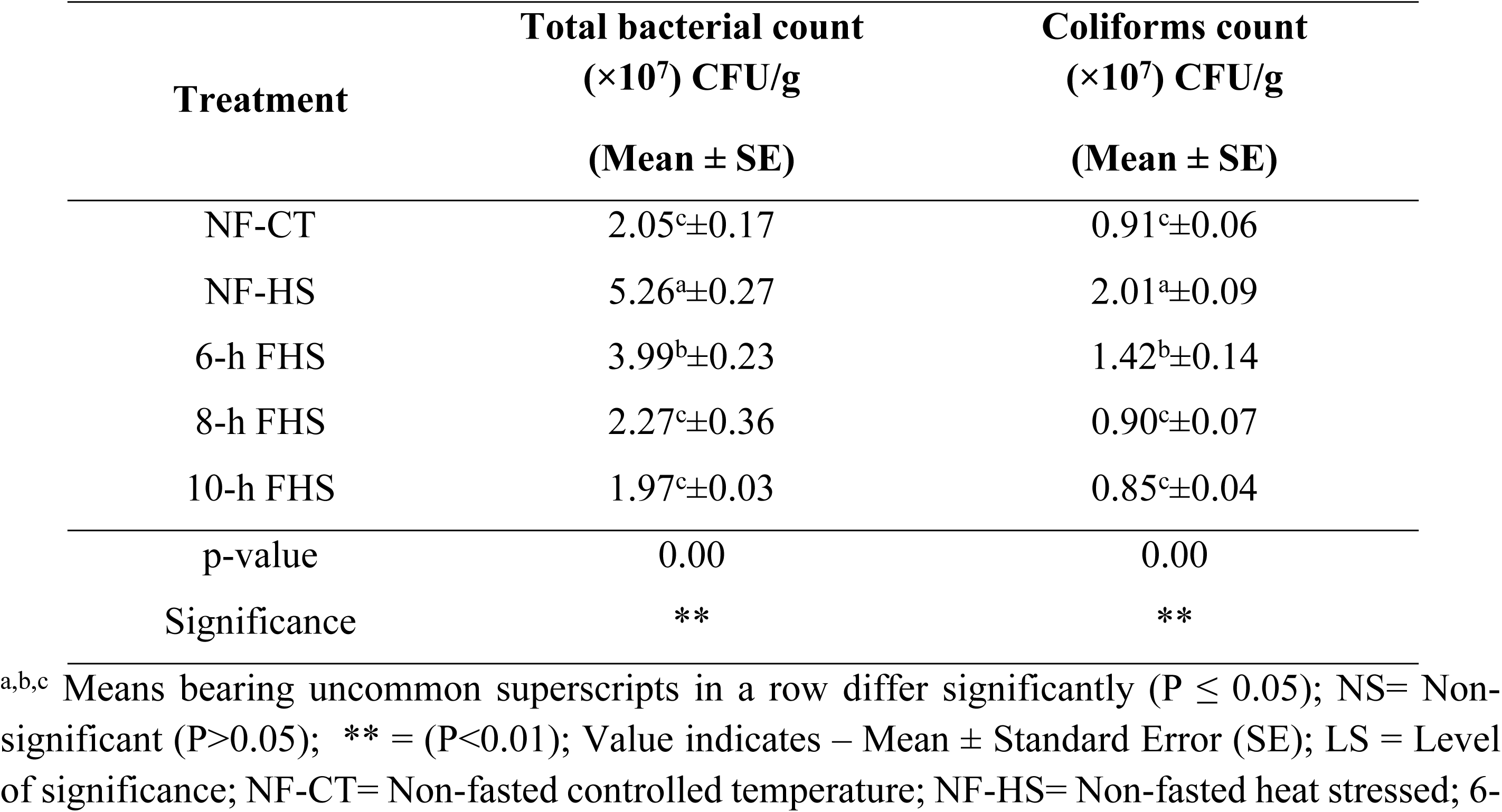
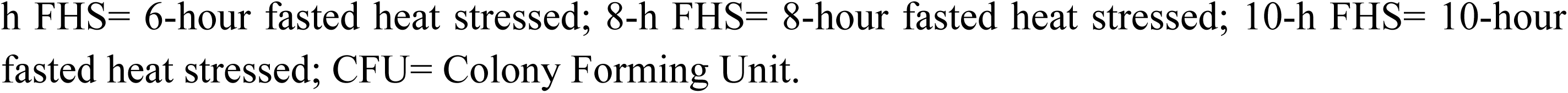
Changes in the cecal microbial profile of broilers exposed to different periodic fasting under controlled and heat-stressed conditions.

| Treatment | Total bacterial count<br>($\times 10^7$) CFU/g | Coliforms count<br>($\times 10^7$) CFU/g |
| --- | --- | --- |
| | (Mean ± SE) | (Mean ± SE) |
| NF-CT | 2.05<sup>c</sup>±0.17 | 0.91<sup>c</sup>±0.06 |
| NF-HS | 5.26<sup>a</sup>±0.27 | 2.01<sup>a</sup>±0.09 |
| 6-h FHS | 3.99<sup>b</sup>±0.23 | 1.42<sup>b</sup>±0.14 |
| 8-h FHS | 2.27<sup>c</sup>±0.36 | 0.90<sup>c</sup>±0.07 |
| 10-h FHS | 1.97<sup>c</sup>±0.03 | 0.85<sup>c</sup>±0.04 |
| p-value | 0.00 | 0.00 |
| Significance | ** | ** |

<sup>a,b,c</sup> Means bearing uncommon superscripts in a row differ significantly ( $P \leq 0.05$ ); NS= Non-significant ( $P > 0.05$ ); \*\* = ( $P < 0.01$ ); Value indicates – Mean ± Standard Error (SE); LS = Level of significance; NF-CT= Non-fasted controlled temperature; NF-HS= Non-fasted heat stressed; 6-
h FHS= 6-hour fasted heat stressed; 8-h FHS= 8-hour fasted heat stressed; 10-h FHS= 10-hour fasted heat stressed; CFU= Colony Forming Unit.

### Benefit-cost analysis

The benefit-cost analysis for the treatment groups is presented in Table 7. The NF-CT group had the highest feed cost per bird (BDT 207.14/-), which was significantly different (p<0.05) from the other groups. The 6-h and 10-h FHS groups had the next highest feed costs, while the NF-HS and 8-h FHS groups had the lowest feed costs (BDT 193.35/- and 195.06/- respectively). Although the overall production cost per bird did not vary considerably, there were notable differences in the total cost per kg of live bird. The NF-HS group had the highest cost (BDT 139.80/-), while the NF-CT group had the lowest cost of production (BDT 121.87/-). Among the treatment groups that were exposed to heat stress, the 8-h FHS group demonstrated the lowest production cost per kg of the live bird, amounting to BDT 131.40/-. The NF-CT group achieved the maximum profit, either per bird or per kg of live bird, compared to the NF-HS group, which had the significantly (p<0.05) lowest profit per bird or per kg of live bird. The benefit-cost ratio (BCR) was statistically higher in the NF-CT group (1.27) and lowest in the NF-HS group (1.11), similar to the profit. Among various FHS groups, the birds that exhibited 8-h fasting under heat stress had a considerably superior benefit-cost ratio (1.18).

**Table 7.** Benefit cost analysis of birds exposed to different fasting period under controlled temperature and heat stressed condition.

| Parameters | NF-CT | NF-HS | 6-h FHS | 8-h FHS | 10-h FHS | p-value | LS |
| --- | --- | --- | --- | --- | --- | --- | --- |
| Feed cost (BDT/bird) | 207.14<sup>a</sup>±2.66 | 193.35<sup>b</sup>±2.52 | 199.17<sup>ab</sup>±4.19 | 195.06<sup>b</sup>±2.47 | 198.38<sup>ab</sup>±2.27 | 0.030 | * |
| Chick cost (BDT/bird) | 19.00 | 19.00 | 19.00 | 19.00 | 19.00 | - | - |
| Bedding, medicine and disinfectant (BD./bird) | 15.00 | 15.00 | 15.00 | 15.00 | 15.00 | - | - |
| Mortality cost (BDT/bird) | 0.00<sup>b</sup>±0.00 | 11.34<sup>a</sup>±4.66 | 3.80<sup>ab</sup>±1.80 | 0.00<sup>b</sup>±0.00 | 0.00<sup>b</sup>±0.00 | 0.029 | * |
| Labor cost (BDT/bird) | 13.00 | 13.00 | 13.00 | 13.00 | 13.00 | - | - |
| Electricity and others (BDT/bird) | 10.00 | 10.00 | 10.00 | 10.00 | 10.00 | - | - |
| Total cost of production (BDT/bird) | 264.14±2.66 | 261.69±4.92 | 259.97±5.17 | 252.07±2.47 | 255.38±2.27 | 0.186 | NS |
| Total cost of production (BDT /kg. body wt.) | 121.87<sup>c</sup>±1.22 | 139.80<sup>a</sup>±3.18 | 135.49<sup>ab</sup>±2.75 | 131.40<sup>b</sup>±2.22 | 137.22<sup>ab</sup>±1.18 | 0.000 | ** |
| Total income (BDT /bird) (Sale price @ BDT 155/kg) | 336.08<sup>a</sup>±4.71 | 290.46<sup>b</sup>±5.73 | 297.58<sup>b</sup>±5.12 | 297.78<sup>b</sup>±6.98 | 288.48<sup>b</sup>±0.95 | 0.000 | ** |
| Profit (BDT/ bird) | 71.94<sup>a</sup>±3.47 | 28.77<sup>c</sup>±6.19 | 37.61<sup>bc</sup>±5.53 | 45.71<sup>b</sup>±5.16 | 33.09<sup>bc</sup>±2.23 | 0.000 | ** |
| Profit (BDT/kg live bird) | 33.13<sup>a</sup>±1.22 | 15.19<sup>c</sup>±3.18 | 19.51<sup>bc</sup>±2.76 | 23.59<sup>b</sup>±2.22 | 17.78<sup>bc</sup>±1.19 | 0.000 | ** |
| Benefit cost ratio (BCR) | 1.27<sup>a</sup>±0.01 | 1.11<sup>c</sup>±0.02 | 1.15<sup>bc</sup>±0.02 | 1.18<sup>b</sup>±0.04 | 1.13<sup>bc</sup>±0.00 | 0.000 | ** |

<sup>abc</sup>Means bearing uncommon superscripts in a row differ significantly ( $P \leq 0.05$ ); NS= Non-significant ( $P > 0.05$ ); \*\* = ( $P < 0.01$ ); \* = ( $P < 0.05$ ); Value indicates – Mean ± Standard Error (SE); LS = Level of significance; NF-CT= Non-fasted controlled temperature; NF-HS= Non-fasted heat stressed; 6-h FHS= 6-hour fasted heat stressed; 8-h FHS= 8-hour fasted heat stressed; 10-h FHS= 10-hour fasted heat stressed; BDT= Bangladeshi taka.

## Discussion

### Temperature humidity index (THI)

The primary measure used to assess the influence of the environment on poultry is the Temperature-Humidity Index (THI) [37], the values of which broilers are classified into four categories: normal (≤ 74), alert (75-78), danger (79-83), and emergency (≥84) [38]. According to Tao and Xin [39], if THI exceeded 75 or 79, it indicated that the birds are facing moderate or severe heat stress. In the present investigation, broilers in the controlled temperature were maintained within a normal THI range (≤ 74) from Day 21 to Day 35. However, birds in the heat-stressed groups experienced extremely elevated THI values ranging from 84 to 89. In general, broilers in tropical and subtropical climates face the challenge of dealing with high THI levels, which are equal to or greater than 84. The study was conducted from July to August, a time when Bangladesh’s natural environment frequently encounters elevated temperatures and extremely high humidity levels. As a result, broilers raised in open-sided houses were exposed to severe environmental threats, and the birds, therefore, they must adapt to such harsh conditions at the cost of their usual growth, physiology, productivity, and reproduction. Thus, the results clearly demonstrated the significance of the current study in showing how different periods-fasting practices minimize the heat stress challenges in experimental broiler chickens.

### Growth performance and mortality

As expected, birds subjected to controlled temperatures (≤26℃) had higher feed consumption and showed better body weight, weight gain, and feed conversion ratio. In contrast, broilers that were exposed to heat stress and subjected to both non-fasting and different fasting schedules exhibited decreased feed intake, resulting in poor growth performance. Several research findings have shown that the broiler chickens exposed to heat stress had reduced feed intake (-14.90% to 28.74%), body weight (-32.6%), and body weight gain (-25.71%); however, an increased in feed conversion ratio (+13.06% to 25.6%) while compared with the birds kept under thermoneutral conditions at 24℃ [40–42]. Broilers under heat stress induce alterations in neuroendocrine activity that especially affect the hypothalamic-pituitary-adrenal axis. This can result in elevated levels of plasma corticosteroid hormones, commonly referred to as ’stress hormones,’ which have the potential to interfere with the feed digestion, nutrient absorption, and utilization in heat-stressed broilers, leading to a decrease in both body weight gain and feed efficiency [42–44].

Previous published reports revealed that the heat stress did not have any impact on the growth performance of commercial broiler chickens during the initial three weeks; however, their FI and BWG were decreased, and FCR was increased at the later part of rearing (from day 22 to 35) [45]. Keeping this point in view, the different fasting approaches in the present study were employed after 21 days and continued till 35 to investigate the impact of various periods of feed withdrawal in minimizing heat stress. In the current study, no significant changes were seen in the body weight and weight gain of broilers among the heat-stressed non-fasted and fasted groups during the first 3 weeks. However, the numerical trend revealed a steady increase in body weight when the fasting period was extended up to 8 hours, followed by a drop-in weight gain with longer fasting durations, resulting in the lowest growth performance in the birds that fasted for 10-h (Table 2). Previous published reports clearly demonstrated that short-term fasting during the last week of the experiment increased the body weight of broilers [26]. The authors further argued and justified that the birds consumed more feed when it was supplied after a fasting period, hence resulting in enhanced growth in starved birds. In general, heat stress hinders the digestive tract’s ability to absorb nutrients, but surprisingly, the birds that undergo fasting have been shown to boost intestinal absorption, leading to improved growth of birds [46].

The overall FCR was decreased along with the increase in fasting duration, reaching its lowest point in the group that fasted for 8-h. However, a protracted fasting period that lasted for 10-h surprisingly resulted in the highest values for FCR, suggesting that the broilers exposed to longer fasting periods exhibited less efficiency in feed utilization. There is scientific evidence suggests that the practice of fasting in broilers under heat stress resulted in several benefits, including enhanced body weight and weight gain, decreased feed consumption, improved feed utilization, and reduced mortality [47–49]. Why the practice of fasting for more than 8-h in broiler exerted poor growth and lower feed utilization remains unclear; however, Shamma et al. [48] have suggested that the extended duration of fasting may lead to a state of ’fasting severity’ in the heat-stressed birds which, in turn, can impede the bird’s ability in effective absorption and utilization of nutrients. Notably, these results are pretty consistent with the findings of the current study.

The broilers in the NF-HS group exhibited a significantly higher mortality compared to the NF-CT group, where no birds died. Even zero mortality was also seen when the birds were subjected to 8-h or 10-h fasting under heat stress. The birds in NF-HS groups had access to continuous feed consumption throughout the day, even during the hottest hours, resulting in increased metabolic heat production. Consequently, there was a significant increase in the bird’s body temperature, causing a disruption in their homeostasis and ultimately resulting in increased mortality. On the contrary, when the birds were fasted for a period of 8- to 10-h, their digestive system remained nearly empty throughout the hottest hours of the day, resulting in minimum stress with almost no mortality in these particular groups, which might be attributed because of less metabolic heat production. Shamma et al. [48] also showed a relationship between the length of fasting and mortality, demonstrating that longer fasting durations were associated with lower mortality. They reported about 7% mortality in the non-fasted groups, but no mortality was recorded up to 28 days after 4-h of fasting or up to 21 days after 6-h. Commercial broilers, even when exposed to a short period of fasting (3-h) during heat stress, showed improved growth performance and decreased mortality as compared to the birds fed *ad-libitum* [49–51] and this could be attributed due to the increased thermal tolerance of the feed-restricted birds [37]. All these results, however, again corroborate with the findings of the current study. Thus, it is likely that the commercial broilers fasted for 8-h in hot and humid conditions are anticipated to have an ameliorating impact on growth performance and survivability under heat stress conditions.

### Impact of fasting on the carcass and dressing yields

The birds with *ad-libitum* access to feed (NF-HS) or were subjected to a longer fasting period (10-h) had the lowest carcass weight and dressing percentage compared to the birds who fasted for 6-h or 8-h. As previously stated, the experimental birds that were maintained in a controlled environment (NF-CT) or subjected to 6-h and 8-h fasting under heat stress exhibited better growth and body weight gain compared to the birds who underwent longer periods of fasting. Thus, it is likely that the birds in these specific groups would have better carcass weight and dressing yield compared to the birds raised either in non-fasting or for longer periods of fasting under heat stress, as observed in the present study. Several published reports also suggest that heat stress exerts a detrimental impact on the growth, carcass, and dressing yields of commercial broilers that undergo *ad libitum* feeding. However, when these birds are subjected to fasting for a standard duration, their growth and carcass quality parameters are significantly boosted [15–17,26,48,52–54].

Along with all other carcass metrics, the overall liver quality was also affected in the broilers fed *ad-libitum* under heat stress (NF-HS groups); however, the restoration of liver weight and overall liver quality has been achieved by 6- or 8-h fasting during the later stages of rearing. Why the fast-growing modern broilers are quite sensible to high environmental temperature (HS) remains unclear; however, it might be possible that the genetic acceleration, nutritional enhancement, and the implementation of current biotechnological tools in today’s broiler operations result in massive growth and make the birds more vulnerable to any adverse environment. Modern broilers subjected to heat stress may cause oxidative damage in the liver tissues, which subsequently disrupts lipid metabolism [55]. Other published reports also postulated significantly lower liver weight in broilers fed *ad libitum* under heat stress [25,56], which could potentially be attributed to the occurrence of oxidative stress within the liver tissue of heat-stressed chickens [57]. Almost similar results of decreased liver weight in broilers of NF-HS groups were depicted by Nisar et al. [58], who further elucidated that the lower liver weight can be attributed to severe tissue damage or oxidation in the liver. The liver is an indispensable organ of the body that primarily cleans the blood by eliminating harmful substances, upholds optimal blood glucose levels, controls blood coagulation, and carries out numerous other essential functions. Hence, commercial broilers reared in open-sided houses under heat stress are recommended for 6- to 8-h fasting in order to keep the liver healthy, standard weight, size, and effectively functional.

Abdominal fat is an inedible component of a carcass that has a detrimental impact on the overall carcass yield and quality. In the current study, the fat percentage of broiler chickens decreased gradually and significantly with the increase in the fasting period, suggesting a notable impact on the accumulation of abdominal fat in birds under heat stress. Hence, the duration of fasting in the heat-stressed broilers is negatively correlated with abdominal fat deposition [49,59], in which the birds under fasting might utilize the stored body fat to maintain their energy requirement. In addition, the abdominal fat reduction in fasted birds under heat stress is somewhat associated with lowering the activity of lipoprotein lipase enzyme (an enzyme that stimulates the release of fatty acids from lipoproteins), which subsequently hinders the accumulation of fat in the abdominal region of the birds [60]. Other published findings also claimed that birds subjected to heat stress exerted lower protein content and higher fat deposition [1,14,15]. The highest abdominal fat in NF-HS groups may be due to the production of the high amount of reactive oxygen species (ROS) in the liver that may alter lipid metabolism and thereby increase the abdominal fat, and reduce the meat yield and meat quality [61].

Finally, the aforementioned explanations have clearly demonstrated that the 8-h fasting under heat stress is a noteworthy strategy to enhance the overall growth, carcass, and dressing yields of broiler chickens. However, this strategy may be impacted adversely when the birds are exposed to *ad-libitum* feeding in highly hot and humid climatic conditions or subjected to prolonged fasting.

### Meat quality of broilers subjected to different fasting regimes

The findings of the current study revealed that breast meat pH was relatively higher in the NF-CT and 8-h FHS groups, while it was lowest in the NF-HS group, suggesting that fasting of birds under heat stress keeps the breast meat pH at a level that almost similar to the birds reared under thermo-neutral comfort environment. In relation to breast meat pH, the highest WHC and lowest CL were found in different period FHS groups. However, poor WHC and highest CL were investigated in the birds of NF-HS. The drip loss however remained unaffected. Corroborating with present findings, various previous reports claimed that subsequent heat stress enhances the level of hydrogen ion [H^+^] and lactic acid mass in the muscle of broilers by accelerating anaerobic glycolysis, which ultimately causes a rapid drop in muscle pH and reducing the WHC of muscle protein [62–64] and thus increasing the drip loss (DL), CL and shear force values in the breast meat of heat-stressed broilers [14–16,65,66]. Therefore, it is evident that a more extended fasting period (8- to 10-h) improved the broiler meat pH and WHC while lowering CL and thus restoring the meat quality that had been compromised by heat stress.

The breast meat color, lightness (L) and yellowness (b*) values remained unchanged but found significantly lowest redness (a*) in the NF-HS group; however, different periods of FHS groups exerted statistically almost similar redness (a*) values like NF-CT. With the agreement of present results, various previous reports also found higher lightness value and lower redness values in the heat-stressed broiler meat [15,16,65,66]. The meat of chickens exposed to heat stress loses the redness value due to the rapid oxidation of myoglobin protein brought on by an abundance of reactive oxygen species [15–16], and also reduces the shelf-life of meat [65–66]. In accordance with high body temperature and lower muscle pH of slaughtered broilers exposed to heat stress increases muscle protein degradation and thus makes the muscle paler and darker [64,67]. On the contrary, Uzum and Toplu [47] found no significant changes in meat color among *ad-libitum* and 8-h fasted groups under heat stress. The hue angle is the indication of discoloration, where the higher values indicate a greater level of discoloration of meat. In the current study, a significant change in hue angle was observed, and a higher hue angle value (77⁰) was found in the breast meat of the NF-HS group. On the other hand, birds of FHS groups depicted hue angle values (52-61⁰) that were almost similar to the birds reared in the thermoneutral zone (NF-CT). Abudabos et al. [68] also postulated numerically higher hue angle value (68⁰) in heat stressed (35℃) birds compared to the birds of thermoneutral control group (61⁰). Again, the saturation index (SI) describes the color intensity of meat, which is found unaffected among the treatment groups in the current study. Although it is statistically insignificant, a numerically lower SI was measured in NF-HS and found higher in 8 to 10-h FHS groups. Davoodi and Ehsani [69] found a SI (13.40) in broilers reared in conventional houses, which are close to controlled temperature groups in the present findings. So, from these findings, it can be concluded that the negative impact of heat stress on broiler breast meat color and quality can be reduced by applying fasting approaches, more specifically for 8-h under hot and humid weather conditions.

### Gut histomorphological changes in the experimental birds

The birds in the NF-CT group exhibited greater villus height (VH), width (VW), and crypt depth (CD) compared to the heat-stressed groups, suggesting that high body temperature in heat-stressed birds leads to the fatal destruction of gut mucosal epithelium cells which finally results in alterations of the gut morphology. Previous research findings demonstrated significant changes in intestinal morphology (VH, VW, VH:CD), and slower daily weight gain in the birds subjected to heat stress [19,70]. Enterocyte proliferation, which occurs in the crypt of the gut, is diminished or inhibited when chickens are subjected to acute heat stress, resulting in lower crypt depth [71–73], which means the limited area for retaining secreting glands in the crypt and so lowering the digestive capabilities of heat-stressed birds. Moreover, Burkholder et al. [18] specifically asserted that the CD exhibited a slight decrease upon the birds’ transition from a higher temperature (30°C) to a lower temperature (23°C) over a 24-h period; however, the mucosa of the duodenum and jejunum demonstrated a rapid change. Gut mucosal damage and morphological alterations may occur in heat-stressed birds as the consequence of oxidative stress and inflammation induced by decreased feed intake and diminished blood supply [74].

In the present study, the levels of VH, VW, CD, and VH:CD ratio in the duodenum, jejunum, and ileum showed a significant increase as the fasting period was extended up to 8-h. However, all these gut health parameters deteriorated with further elongation of the fasting time in the birds under heat-stressed conditions. Almost similar results were depicted by Shamma et al. [48] and Azouz [75]. These increased villi length and width in the intestine provide more surface area for the absorption of nutrients, thereby enhancing improved nutrient digestibility and utilization. Consequently, this leads to improved growth and feed conversion ratio (FCR) in feed-restricted groups, even when the birds are subjected to heat stress conditions, in comparison to *ad-libitum* feeding [76–78]. Thompson and Applegate [79] also showed that feed restriction increased the jejunal villus height of chickens at six weeks of age, which could be attributed to an adaptive strategy to maximize nutrient uptake during feeding. The findings mentioned above, in conjunction with the outcomes of the present investigation, unequivocally indicate that fast-growing modern broilers, when exposed to 8-h fasting in hot-humid climatic conditions, exhibited notable enhancements in their overall gut health, improved nutrient absorption, and positively influenced their growth performance.

### Impact of different fasting regimes on cecal microbial profile

In the current study, the bacterial and coliform counts in the cecal content of broilers in the NF-HS groups were found to be considerably higher compared to birds of NF-CT or subjected to different periods of fasting under heat stress. The broiler chickens reared at high ambient temperatures significantly increased the total cecal bacterial counts [80]; even the excessive heat stress promoted the proliferation of harmful pathogenic bacteria such as *E. coli* and *Salmonella*, as well as total aerobic bacteria, rather than the beneficial microflora found in the cecum [81]. Several published reports have shown that the antioxidant defense mechanism of heat-stressed broiler chickens reduces lipid peroxidation, leading to alterations in the bacterial communities in the intestine. This alteration is characterized by an increase in harmful bacteria (*Coliforms* and *Clostridium*) and/or a decrease in beneficial bacteria (*Lactobacillus* and *Bifidobacterium*), which consequently has a detrimental impact on gut health, digestion, and nutrients absorption [19,82–84]. Typically, broilers experiencing heat stress lose their intestinal integrity, leading to an elevation in the permeability and penetration of harmful bacteria such as *Salmonella spp* [85], which is considered one of the major causes of high mortality in heat-stressed broilers [44]. The present investigation revealed relatively higher mortality among broilers subjected to heat stress while receiving *ad libitum* feeding, which might be because of the infiltration of harmful bacteria in the intestine. The results of the present study have also shown a substantial decrease in the overall bacterial count, and *Coliform* count in the caecum of broilers subjected to heat stress, along with an increase in the length of fasting. Shafiei et al. [56] suggested and provided evidence that the inadequate production of pancreatic juice in the gastrointestinal tract of broiler chickens that underwent prolonged fasting created a favored pH level that promoted the proliferation of *Lactobacillus* bacteria. Consequently, the prevailing circumstances fostered a conducive environment for the proliferation of beneficial bacteria but suppressed the number of pathogenic bacteria within the gastrointestinal tract. All of these aforementioned research findings corroborate the result outcomes of the current investigation.

### Benefit-cost analysis

As anticipated, the birds housed in a controlled environment (NF-CT group) exhibited a higher overall feed cost due to their increased amount of feed consumption under a comfortable thermal environment. However, broilers in this group showed zero mortality and better feed utilization, leading to the lowest production cost per kg or per piece of live bird that eventually yields a higher profit or benefit-cost ratio (BCR). In contrast, when experimental birds are provided with constant access to feed, even in the warmest part of the day, it leads to significant energy depletion as a consequence of persistent panting under heat stress. This, in turn, leads to reduced feed intake, impaired nutrient utilization, and elevated mortality rates. The combination of extreme ambient temperatures and the metabolic heat generated by broilers themselves in a hot and humid environment likely disrupts the usual physiological processes of homeostasis in this particular type of bird. Collectively, these factors are attributed to elevated production costs and the lowest profit, accompanied by minimal BCR in comparison to other treatment groups. Remarkably, when heat-stressed birds were subjected to fasting, they exhibited slightly better feed utilization, resulting in reduced mortality than those fed *ad libitum*. Consequently, the broiler chickens those were subjected to different fasting durations, resulting in lower production costs and increased profitability with moderate BCR values. However, when comparing different fasting groups, it was seen that birds fasted for 8-h generated considerably higher levels of profit and BCR compared to both fasted and non-fasted heat-challenged groups.

Several published articles have provided evidence supporting the findings of the current study, demonstrating that feed restrictions, both in terms of quantity and time duration, yield a net economic benefit when compared to full feeding of broilers under heat stress [75,86,87].

## Conclusion

Based on the results obtained from the current study, it can be concluded that boiler chickens exposed to 8-h of fasting during the hottest period of the day in summer is an effective way to mitigate the effect of heat stress under hot and humid climatic conditions of the tropical and sub-tropical region including Bangladesh. This 8-h fasting regimen assures that birds grow and produce at their best as possible with little to no mortality loss by improving gut health and the microbial ecosystem, even in the face of heat stress. Consequently, it is readily acceptable and adaptable for broiler farmers of various scales, including those operating small, medium, or even large farms. Part II of our study will focus on birds’ behavioral, physiological, hematological, and immunological states employing the same fasting regimens under heat stress.

## Acknowledgements

The Department of Poultry Science and Department of Microbiology and Hygiene of Bangladesh Agricultural University provided logistical support and other laboratory facilities, for which the authors are highly grateful. The funding for this research was granted by the Ministry of Science and Technology (MoST) of Bangladesh (Projrct No: 2021/121/MoST).

## Supporting information

**S1 Fig. S1 Fig. Histomorphological changes in the small intestine of broiler chickens among different treatment groups (image: 4x).**

## Notes

### Competing Interest Statement

The authors have declared no competing interest.

## References

1. Lara LJ, Rostagno MH. Impact of heat stress on poultry production. Animals. 2013; 3(2):356–69. 10.3390%2Fani3020356

2. IFRC 2024: Bangladesh/Heatwave-DREF Operation Appeal: MDRBD034. Available from: https://reliefweb.int/report/bangladesh/bangladesh-heatwave-dref-operation-appeal-mdrbd034

3. Tellez Jr G, Tellez-Isaias G, Dridi S. Heat stress and gut health in broilers: role of tight junction proteins. Advances in Food Technology and Nutritional Sciences. 2017; 3:e1–4. 10.17140/AFTNSOJ-3-e010

4. Kumar M, Ratwan P, Dahiya SP, Nehra AK. Climate change and heat stress: Impact on production, reproduction and growth performance of poultry and its mitigation using genetic strategies. Journal of thermal biology. 2021; 97:102867. 10.1016/j.jtherbio.2021.102867

5. Sukanta H, Monjurul H, Sushanta G. Poultry farmers in a bind as chickens succumb to heatwave. The Daily Star. 2024 April 26. Available from: https://www.thedailystar.net/business/economy/news/poultry-farmers-bind-chickens-succumb-heatwave-3595436

6. Rokon U. Heatwave may cut meat, milk & egg production by 25%. The Business Post. 2024 Apr 27. Available from: https://businesspostbd.com/national/heatwave-may-cut-meat-milk-egg-production-by-25#google_vignette

7. Charles, D. R. Responses to the thermal environment. In D. A. Charles, & A. W. Walker (Eds.), Poultry environment problems, a guide to solutions. Nottingham University Press. 2002; Pages 1–16.

8. Olanrewaju HA, Purswell JL, Collier SD, Branton SL. Effect of ambient temperature and light intensity on physiological reactions of heavy broiler chickens. Poultry Science. 2010; 89(12):2668–77. 10.3382/ps.2010-00806

9. Oke OE, Alo ET, Oke FO, Oyebamiji YA, Ijaiya MA, Odefemi MA, Kazeem RY, Soyode AA, Aruwajoye OM, Ojo RT, Adeosun SM. Early age thermal manipulation on the performance and physiological response of broiler chickens under hot humid tropical climate. Journal of Thermal Biology. 2020; 88:102517. 10.1016/j.jtherbio.2020.102517

10. Zhang C, Zhao XH, Yang L, Chen XY, Jiang RS, Jin SH, Geng ZY. Resveratrol alleviates heat stress-induced impairment of intestinal morphology, microflora, and barrier integrity in broilers. Poultry science. 2017; 96(12):4325–32. 10.3382/ps/pex266

11. Mohanaselvan A, Bhaskar E. Mortality from non-exertional heat stroke still high in India. The international journal of occupational and environmental medicine. 2014; 5(4):222.

12. Khan RU, Naz S, Nikousefat Z, Tufarelli V, Javdani M, Rana N, Laudadio V. Effect of vitamin E in heat-stressed poultry. World’s poultry science journal. 2011; 67(3):469–78. 10.1017/S0043933911000511

13. Van Goor A, Bolek KJ, Ashwell CM, Persia ME, Rothschild MF, Schmidt CJ, Lamont SJ. Identification of quantitative trait loci for body temperature, body weight, breast yield, and digestibility in an advanced intercross line of chickens under heat stress. Genetics Selection Evolution. 2015; 47:1–3. 10.1186/s12711-015-0176-7

14. Lu Q, Wen J, Zhang H. Effect of chronic heat exposure on fat deposition and meat quality in two genetic types of chicken. Poultry science. 2007; 86(6):1059–64. 10.1093/ps/86.6.1059

15. Zhang ZY, Jia GQ, Zuo JJ, Zhang Y, Lei J, Ren L, Feng DY. Effects of constant and cyclic heat stress on muscle metabolism and meat quality of broiler breast fillet and thigh meat. Poultry science. 2012a; 91(11):2931–7. 10.3382/ps.2012-02255

16. Zhang M, Zou XT, Li H, Dong XY, Zhao W. Effect of dietary γ-aminobutyric acid on laying performance, egg quality, immune activity and endocrine hormone in heat-stressed Roman hens. Animal Science Journal. 2012b; 83(2):141–7. 10.1111/j.1740-0929.2011.00939.x

17. Zeferino CP, Komiyama CM, Pelícia VC, Fascina VB, Aoyagi MM, Coutinho LL, Sartori JR, Moura AS. Carcass and meat quality traits of chickens fed diets concurrently supplemented with vitamins C and E under constant heat stress. Animal. 2016; 10(1):163–71. 10.1017/S1751731115001998

18. Burkholder KM, Thompson KL, Einstein ME, Applegate TJ, Patterson JA. Influence of stressors on normal intestinal microbiota, intestinal morphology, and susceptibility to Salmonella enteritidis colonization in broilers. Poultry science. 2008; 87(9):1734–41. 10.3382/ps.2008-00107

19. Song J, Xiao K, Ke YL, Jiao LF, Hu CH, Diao QY, Shi B, Zou XT. Effect of a probiotic mixture on intestinal microflora, morphology, and barrier integrity of broilers subjected to heat stress. Poultry science. 2014; 93(3):581–8. 10.3382/ps.2013-03455

20. Santos RR, Awati A, Roubos-van den Hil PJ, Tersteeg-Zijderveld MH, Koolmees PA, Fink-Gremmels J. Quantitative histo-morphometric analysis of heat-stress-related damage in the small intestines of broiler chickens. Avian Pathology. 2015; 44(1):19–22. 10.1080/03079457.2014.988122

21. He X, Lu Z, Ma B, Zhang L, Li J, Jiang Y, Zhou G, Gao F. Effects of chronic heat exposure on growth performance, intestinal epithelial histology, appetite-related hormones and genes expression in broilers. Journal of the Science of Food and Agriculture. 2018; 98(12):4471–8. 10.1002/jsfa.8971

22. Sohail MU, Hume ME, Byrd JA, Nisbet DJ, Shabbir MZ, Ijaz A, Rehman H. Molecular analysis of the caecal and tracheal microbiome of heat-stressed broilers supplemented with prebiotic and probiotic. Avian pathology. 2015; 44(2):67–74. 10.1080/03079457.2015.1004622

23. Wang XJ, Feng JH, Zhang MH, Li XM, Ma DD, Chang SS. Effects of high ambient temperature on the community structure and composition of ileal microbiome of broilers. Poultry Science. 2018; 97(6):2153–8. 10.3382/ps/pey032

24. Zhu L, Liao R, Wu N, Zhu G, Yang C. Heat stress mediates changes in fecal microbiome and functional pathways of laying hens. Applied microbiology and biotechnology. 2019; 103:461–72. 10.1007/s00253-018-9465-8

25. Azis A, Afriani A. Effect of feeding time restriction during the growing period on growth performance of broiler chickens. Asian Journal of Poultry Science. 2017; 11(2):70–4. 10.3923/ajpsaj.2017.70.74

26. Özkan S, Akbaş Y, Altan Ö, Altan A, Ayhan V, Özkan K. The effect of short-term fasting on performance traits and rectal temperature of broilers during the summer season. British poultry science. 2003; 44(1):88–95. 10.1080/0007166031000085292

27. Susbilla JP, Tarvid I, Gow CB, Frankel TL. Quantitative feed restriction or meal-feeding of broiler chicks alter functional development of enzymes for protein digestion. British poultry science. 2003; 44(5):698–709. 10.1080/00071660310001643679

28. Zhan XA, Wang M, Ren H, Zhao RQ, Li JX, Tan ZL. Effect of early feed restriction on metabolic programming and compensatory growth in broiler chickens. Poultry Science. 2007; 86(4):654–60. 10.1093/ps/86.4.654

29. Lindholm C, Altimiras J. Physiological and behavioural effects of intermittent fasting vs daily caloric restriction in meat-type poultry. Animal. 2023; 17(6):100849. 10.1016/j.animal.2023.100849

30. Zhou WT, Fujita M, Ito T, Yamamoto S. Effects of early heat exposure on thermoregulatory responses and blood viscosity of broilers prior to marketing. British Poultry Science. 1997; 38(3):301–6. 10.1080/00071669708417991

31. Leeson S. Nutritional considerations of poultry during heat stress. World’s Poultry Science Journal. 1986; 42(1):69–81. 10.1079/WPS19860007

32. Meat Color Measurement Guideline. American Meat Science Association. 2012. Available from: https://meatscience.org/docs/default-source/publications-resources/hottopics/2012_12_meat_clr_guide.pdf?sfvrsn=d818b8b3_0

33. Afrin A, Ahmed T, Lahiry A, Rahman S, Dey B, Hashem MA, Das SC. Profitability and meat quality of fast-, medium-and slow-growing meat-type chicken genotypes as affected by growth and length of rearing. Saudi Journal of Biological Sciences. 2024; 31(8):104025. 10.1016/j.sjbs.2024.104025

34. Marai IF, Ayyat MS, Abd El-Monem UM. Growth performance and reproductive traits at first parity of New Zealand White female rabbits as affected by heat stress and its alleviation under Egyptian conditions. Tropical animal health and production. 2001; 33:451–62. 10.1023/a:1012772311177

35. Hahn GL, Gaughan JB, Mader TL, Eigenberg RA. Thermal indices and their applications for livestock environments. InLivestock energetics and thermal environment management 2009 (pp. 113–130). American Society of Agricultural and Biological Engineers. Available from: https://www.google.com/search?client=firefox-bd&q=doi%3A10.13031%2F2013.28298

36. SPSS 2013. Statistical Computer Package Program 20.00 (SPSS Inc. Chicago, USA).

37. Lin H, Jiao HC, Buyse J, Decuypere E. Strategies for preventing heat stress in poultry. World’s Poultry Science Journal. 2006; 62(1):71–86. 10.1079/WPS200585

38. LCI. Patterns of transit losses. Omaha, Neb.: Livestock Conservation, Inc. 1970.

39. Tao X, Xin H. Temperature-humidity-velocity index for market-size broilers. In2003 ASAE Annual Meeting 2003 (p. 1). American Society of Agricultural and Biological Engineers.

40. Sohail MU, Hume ME, Byrd JA, Nisbet DJ, Ijaz A, Sohail A, Shabbir MZ, Rehman H. Effect of supplementation of prebiotic mannan-oligosaccharides and probiotic mixture on growth performance of broilers subjected to chronic heat stress. Poultry science. 2012; 91(9):2235–40. 10.3382/ps.2012-02182

41. Sun X, Zhang H, Sheikhahmadi A, Wang Y, Jiao H, Lin H, Song Z. Effects of heat stress on the gene expression of nutrient transporters in the jejunum of broiler chickens (Gallus gallus domesticus). International journal of biometeorology. 2015; 59:127–35. 10.1007/s00484-014-0829-1

42. Olfati A, Mojtahedin A, Sadeghi T, Akbari M, Martínez-Pastor F. Comparison of growth performance and immune responses of broiler chicks reared under heat stress, cold stress and thermoneutral conditions. Spanish Journal of Agricultural Research. 2018; 16(2):e0505-. 10.5424/sjar/2018162-12753

43. Quinteiro-Filho WM, Gomes AV, Pinheiro ML, Ribeiro A, Ferraz-de-Paula V, Astolfi-Ferreira CS, Ferreira AJ, Palermo-Neto J. Heat stress impairs performance and induces intestinal inflammation in broiler chickens infected with Salmonella Enteritidis. Avian Pathology. 2012; 41(5):421–7. 10.1080/03079457.2012.709315

44. Liu L, Ren M, Ren K, Jin Y, Yan M. Heat stress impacts on broiler performance: a systematic review and meta-analysis. Poultry Science. 2020; 99(11):6205–11. 10.1016/j.psj.2020.08.019

45. Awad EA, Najaa M, Zulaikha ZA, Zulkifli I, Soleimani AF. Effects of heat stress on growth performance, selected physiological and immunological parameters, caecal microflora, and meat quality in two broiler strains. Asian-Australasian journal of animal sciences. 2020; 33(5):778–787. 10.5713/ajas.19.0208

46. Pinheiro DF, Cruz VC, Sartori JR, Paulino MV. Effect of early feed restriction and enzyme supplementation on digestive enzyme activities in broilers. Poultry science. 2004; 83(9):1544–50. 10.1093/ps/83.9.1544

47. Uzum MH, Toplu HO. Effects of stocking density and feed restriction on performance, carcass, meat quality characteristics and some stress parameters in broilers under heat stress. Revue de Médecine Vétérinaire. 2013; 164(12):546–54.

48. Shamma TA, Khalifa HH, El-Shafei AA, Abo-Gabal MS. Mitigating heat stress in broilers: 1-effect of feed restriction and early heat acclimation on productive performance. Middle East Journal of Applied Sciences. 2014; 4(4):967–82.

49. Mohamed AS, Lozovskiy AR, Ali AM. Strategies to combat the deleterious impacts of heat stress through feed restrictions and dietary supplementation (vitamins, minerals) in broilers. Journal of the Indonesian Tropical Animal Agriculture. 2019; 44:155–66. 10.14710/jitaa.44.2.155-166

50. Al-Aqil A, Zulkifli I, Sazili AQ, Omar AR, Rajion MA. The effects of the hot, humid tropical climate and early age feed restriction on stress and fear responses, and performance in broiler chickens. Asian-Australasian Journal of Animal Sciences. 2009; 22(11):1581–6. 10.5713/ajas.2009.90021

51. Teyssier JR, Brugaletta G, Sirri F, Dridi S, Rochell SJ. A review of heat stress in chickens. Part II: Insights into protein and energy utilization and feeding. Frontiers in Physiology. 2022; 13:943612. 10.3389/fphys.2022.943612

52. David LS, Subalini E. Effects of Feed restriction on the growth performance, organ size and carcass characteristics of Broiler chickens. Scholars Journal of Agriculture and Veterinary Sciences. 2015; 2(2A):108–11.

53. Zeferino CP, Komiyama CM, Fernandes S, Sartori JR, Teixeira PS, Moura AS. Carcass and meat quality traits of rabbits under heat stress. Animal. 2013; 7(3):518–23. 10.1017/S1751731112001838

54. Jahejo AR, Rajput N, Rajput NM, Leghari IH, Kaleri RR, Mangi RA, Sheikh MK, Pirzado MZ. Effects of heat stress on the performance of Hubbard broiler chicken. Cells, Animal and Therapeutics. 2016; 2(1):1–5.

55. Emami NK, Jung U, Voy B, Dridi S. Radical response: effects of heat stress-induced oxidative stress on lipid metabolism in the avian liver. Antioxidants. 2020; 10(1):35. 10.3390/antiox10010035

56. Shafiei A, Khavarinezhad S, Javandel F, Nosrati M, Seidavi A, Diarra SS. Effects of duration of early feed withdrawal and re-feeding on growth, carcass traits, plasma constituents and intestinal microflora of broiler chickens. Journal of Applied Animal Research. 2018; 46(1):1358–62. 10.1080/09712119.2018.1509004

57. Zhao Y, Zhuang Y, Shi Y, Xu Z, Zhou C, Guo L, Liu P, Wu C, Hu R, Hu G, Guo X. Effects of N-acetyl-l-cysteine on heat stress-induced oxidative stress and inflammation in the hypothalamus of hens. Journal of Thermal Biology. 2021; 98:102927. 10.1016/j.jtherbio.2021.102927

58. Nisar H, Sharif M, Rahman MA, Rehman S, Kamboh AA, Saeed M. Effects of dietary supplementations of synbiotics on growth performance, carcass characteristics and nutrient digestibility of broiler chicken. Brazilian Journal of Poultry Science. 2021; 23(02):eRBCA-2020. 10.1590/1806-9061-2020-1388

59. Saleh A, Alkhamisi AR, Abdul Rahman A, Albayomy MA, Draz M. Influence of Feed Withdrawal Period on Growth Performance of Broiler Chickens under High Ambient Temperature. Journal of Sustainable Agricultural Sciences. 2019; 45(2):59–65. 10.21608/jsas.2019.10669.1136

60. Ghazanfari S, Kermanshahi H, Nassiry MR, Golian A, Moussavi AR, Salehi A. Effect of feed restriction and different energy and protein levels of the diet on growth performance and growth hormone in broiler chickens. Journal of Biological Sciences. 2010; 10(1):25–30. Available from: https://scialert.net/fulltext/fulltextpdf.php?pdf=ansinet/jbs/2010/25-30.pdf

61. Fouad AM, Chen W, Ruan D, Wang S, Xia WG, Zheng CT. Impact of heat stress on meat, egg quality, immunity and fertility in poultry and nutritional factors that overcome these effects: A review. International Journal of Poultry Science. 2016; 15(3):81. 10.3923/ijps.2016.81.95

62. Van Laack RL, Liu CH, Smith MO, Loveday HD. Characteristics of pale, soft, exudative broiler breast meat. Poultry Science. 2000; 79(7):1057–61. 10.1093/ps/79.7.1057

63. Strasburg GM, Chiang W. Pale, soft, exudative turkey—The role of ryanodine receptor variation in meat quality. Poultry Science. 2009; 88(7):1497–505. 10.3382/ps.2009-00181

64. Nawaz AH, Amoah K, Leng QY, Zheng JH, Zhang WL, Zhang L. Poultry response to heat stress: Its physiological, metabolic, and genetic implications on meat production and quality including strategies to improve broiler production in a warming world. Frontiers in veterinary science. 2021; 8:699081. 10.3389/fvets.2021.699081

65. Wang RR, Pan XJ, Peng ZQ. Effects of heat exposure on muscle oxidation and protein functionalities of pectoralis majors in broilers. Poultry Science. 2009; 88(5):1078–84. 10.3382/ps.2008-00094

66. Hao Y, Gu XH. Effects of heat shock protein 90 expression on pectoralis major oxidation in broilers exposed to acute heat stress. Poultry Science. 2014; 93(11):2709–17. 10.3382/ps.2014-03993

67. Wilhelm AE, Maganhini MB, Hernández-Blazquez FJ, Ida EI, Shimokomaki M. Protease activity and the ultrastructure of broiler chicken PSE (pale, soft, exudative) meat. Food chemistry. 2010; 119(3):1201–4. 10.1016/j.foodchem.2009.08.034

68. Abudabos AM, Suliman GM, Al-Owaimer AN, Sulaiman AR, Alharthi AS. Effects of nano emulsified vegetable oil and betaine on growth traits and meat characteristics of broiler chickens reared under cyclic heat stress. Animals. 2021; 11(7):1911. 10.3390/ani11071911

69. Davoodi P, Ehsani A. Characteristics of carcass traits and meat quality of broiler chickens reared under conventional and free-range systems. Journal of World’s Poultry Research. 2020; 10(4):623–30. 10.36380/jwpr.2020.71

70. He J, He Y, Pan D, Cao J, Sun Y, Zeng X. Associations of gut microbiota with heat stress-induced changes of growth, fat deposition, intestinal morphology, and antioxidant capacity in ducks. Frontiers in Microbiology. 2019; 10:903. 10.3389/fmicb.2019.00903

71. Marchini CF, Café MB, Araújo EG, Nascimento MR. Physiology, cell dynamics of small intestinal mucosa, and performance of broiler chickens under heat stress: a review. Revista Colombiana de Ciencias Pecuarias. 2016; 29(3):159–68.

72. Marchini CF, Silva PL, Nascimento MR, Beletti ME, Silva NM, Guimarães EC. Body weight, intestinal morphometry and cell proliferation of broiler chickens submitted to cyclic heat stress. Int. J. Poult. Sci. 2011; 10(6):455–60.

73. Elnesr S, Abdel-Azim A. The impact of heat stress on the gastrointestinal tract integrity of poultry. Labyrinth: Fayoum Journal of Science and Interdisciplinary Studies. 2023;.2:82–90. 10.21608/ifjsis.2023.220540.1031

74. Rostagno MH. Effects of heat stress on the gut health of poultry. Journal of animal science. 2020; 98(4):skaa090. 10.1093/jas/skaa090

75. Azouz HMM., Gadelrab SS, EL-Komy HM. Effects of late feed restriction on growth performance and intestinal villi parameters of broiler chicks under summer conditions. Egyptian Poultry Science Journal. 2019; 39(4):913–34. 10.21608/epsj.2019.67514

76. Yamauchi K, Tarachai P. Changes in intestinal villi, cell area and intracellular autophagic vacuoles related to intestinal function in chickens. British Poultry Science. 2000; 41(4):416–23. 10.1080/00071660050194902

77. Onderci M, Sahin N, Sahin K, Cikim G, Aydin A, Ozercan I, Aydin S. Efficacy of supplementation of α-amylase-producing bacterial culture on the performance, nutrient use, and gut morphology of broiler chickens fed a corn-based diet. Poultry Science. 2006; 85(3):505–10. 10.1093/ps/85.3.505

78. Buwjoom T, Yamauchi K, Erikawa T, Goto H. Histological intestinal alterations in chickens fed low protein diet. Journal of Animal Physiology and Animal Nutrition. 2010; 94(3):354–61. 10.1111/j.1439-0396.2008.00915.x

79. Thompson KL, Applegate TJ. Feed withdrawal alters small-intestinal morphology and mucus of broilers. Poultry science. 2006; 85(9):1535–40. 10.1093/ps/85.9.1535

80. Kammon A, Alzentani S, Tarhuni O, Asheg A. Research Article Effect of Some Organic Acids on Body Weight, Immunity and Cecal Bacterial Count of Chicken during Heat Stress International Journal of Poultry Science. 2019; 18(6):293–300. 10.3923/ijps.2019.293.300

81. Park SO, Hwangbo J, Ryu CM, Park BS, Chae HS, Choi HC, Kang HK, Seo OS, Choi YH. Effects of Extreme Heat Stress on Growth Performance, Lymphoid Organ, IgG and Cecum Microflora of Broiler Chickens. International Journal of Agriculture & Biology. 2013; 15(6) 1204–1208. https://www.fspublishers.org/published_papers/75376_..pdf

82. Song J, Jiao LF, Xiao K, Luan ZS, Hu CH, Shi B, Zhan XA. Cello-oligosaccharide ameliorates heat stress-induced impairment of intestinal microflora, morphology and barrier integrity in broilers. Animal feed science and technology. 2013; 185(3-4):175–81. 10.1016/j.anifeedsci.2013.08.001

83. Al-Fataftah AR, Abdelqader A. Effects of dietary Bacillus subtilis on heat-stressed broilers performance, intestinal morphology and microflora composition. Animal feed science and technology. 2014; 198:279–85. 10.1016/j.anifeedsci.2014.10.012

84. Abdelqader A, Al-Fataftah AR. Effect of dietary butyric acid on performance, intestinal morphology, microflora composition and intestinal recovery of heat-stressed broilers. Livestock Science. 2016; 183:78–83. 10.1016/j.livsci.2015.11.026

85. Alhenaky A, Abdelqader A, Abuajamieh M, Al-Fataftah AR. The effect of heat stress on intestinal integrity and Salmonella invasion in broiler birds. Journal of thermal biology. 2017; 70:9–14. 10.1016/j.jtherbio.2017.10.015

86. Novel DJ, Ng’Ambi JW, Norris D, Mbajiorgu CA. Effect of different feed restriction regimes during the starter stage on productivity and carcass characteristics of male and female Ross 308 broiler chickens. International Journal of Poultry Science. 2009; 8(1):35–39. 10.3923/ijps.2009.35.39

87. Farghly MF, Makled MN. Application of intermittent feeding and flash lighting regimens in broiler chickens management. Egyptian Journal of Nutrition and Feeds. 2015; 18(2):261–76. 10.21608/ejnf.2015.105816

